# Recombinant Photo-tagging Enables Fluorescent Labelling of Biomolecules and Visualization of Liquid–Liquid Phase Separation

**DOI:** 10.64898/2026.01.13.699169

**Authors:** Krishna Agrawal, P S Sivaprasad, Reman Kumar Singh, Yashwant Kumar, Karthik Pushpavanam

## Abstract

Liquid–liquid phase separation (LLPS) of biomolecules has emerged as a fundamental principle governing biological systems. A key aspect to understand LLPS is the ability to visualize these coacervates, which requires labelling strategies. Fluorescent labelling of biomolecules remains one of the widely adopted techniques involving either small-molecule chemical conjugation or recombinant expression alongside a fluorescent protein. These small-molecule fluorophores are typically employed in excess, which further necessitates additional downstream processing steps, including separation of the conjugated molecules from the unreacted fluorophores. On the other hand, recombinantly appended fluorescent protein affects the functioning of the labelled biomolecule owing to its large size. There is still a need to develop a labelling technique with small molecular weight fluorophores that is also genetically encodable. We previously discovered that the C-terminal peptide fragment (CTPF) released during photo-exposure of a green-to-red photoconvertible fluorescent protein (PhoCl1) displayed red fluorescence. Notably, this chromophore displays broad excitation and emission wavelength profiles, rendering it robust under multiple excitation wavelengths. We leverage this phenomenon to achieve post-expression fluorescent labelling by genetically fusing PhoCl1 to the target peptide or protein to be labelled (TPTL). We successfully demonstrated this approach of conferring fluorescence to silica binding peptide, riboflavin kinase enzyme, TEV protease, Lanthanide Binding Peptide and maltose binding protein, all of which exhibited red fluorescence upon photocleavage. Finally, we demonstrate the applicability of this probe to visualize LLPS of CTPF tagged elastin-like polypeptide in the presence of polymeric crowder. We anticipate that this strategy for inducing red fluorescence in non-fluorescent biomolecules via a photocleavable protein will open new avenues for minimally perturbative fluorescent labelling.

## INTRODUCTION

Intracellular partitioning enables cells to perform multiple biochemical processes simultaneously by spatially organizing distinct functions.^1^ While classical compartmentalization relies on membrane-bound organelles such as the nucleus, mitochondria, and Golgi apparatus,^1^ liquid–liquid phase separation (LLPS) provides a membrane-free mechanism for cellular organization through the formation of dynamic biomolecular condensates. LLPS of proteins have been demonstrated to play a crucial role in various cellular processes, including transcription, immune signaling, and autophagy, and is closely linked to human health and diseases.^2^ Despite its importance, visualizing these LLPS remains challenging using conventional optical techniques such as brightfield microscopy. Brightfield imaging relies on differences in light absorption and refractive index and therefore provides limited contrast for transparent, compositionally similar, and dynamic phase-separated assemblies.^3^ Consequently, effective visualization of phase-separated condensates requires appropriate labelling strategies to enable selective and high-contrast observation using optical microscopy platforms.^3^

Proteins and peptides exhibit a diverse range of functions, from self-assembly to catalysis, owing to their ability to adopt unique structures through a wide variety of natural and unnatural amino acid building blocks.^4,5^ Labeling biomolecules is essential in these applications, as it allows for precise tracking of their behavior, interactions, and functionality within complex systems. This detailed monitoring enables the optimization of biomolecule-based technologies for fields including but not limited to bioelectronics, regenerative medicine and catalysis.^6^ There exists a diverse array of techniques available to visualize and track proteins, ranging from radioactive labeling, isotope markers, and electrochemical probes.^6^ Even with this wide range of available techniques, fluorescent labeling remains one of the widely adopted techniques.^7^ Fluorescent labeling offers distinct advantages over its counterpart (1) Non-destructive nature as compared to radioactive labeling techniques, (2) Non-invasive mode of detection as compared to exploiting electrochemical probes and (3) Low concentrations of labeled material can be quantified as compared to isotope labeling.^8,9^ Taken together, these make fluorescent labeling versatile and sensitive for studying biological systems.

A range of fluorescent labeling techniques have been developed over the years, primarily involving chemical-based conjugation methods that covalently attach fluorophores to specific reactive groups on proteins.^10,11^ The N-hydroxysuccinimide (NHS) ester reacts with the amine to form a stable amide bond, effectively attaching the fluorophore to the protein.^12,13^ Another widely used method involves thiol-reactive chemistry, where maleimide or iodoacetamide-based dyes are conjugated to cysteine residues, forming a thioether bond.^14,15^ These small molecule fluorophores are typically employed in excess, which further necessitates additional downstream processing steps, including separation of the conjugated molecules from the unreacted fluorophores. To overcome this, fluorescent proteins have been recombinantly appended, but its interaction with the protein of interest (POI) likely affects the functioning of the labelled protein due to its large size.^16^ Although these approaches have been widely successful, there is still a need to develop a recombinant technique to label proteins with small fluorophores (<2000 Da) which are genetically encodable.

PhoCl1, a circularly permuted variant of β-barrel fluorescent protein mMaple, is designed to confer photocleavability upon 405 nm light exposure.^17^ The chromophore, along with the 9-amino acid C-terminal loop (NRVFTKYPR) within the β-barrel core of PhoCl1, detaches from the protein backbone chain upon 405 nm irradiation. For brevity, we will refer to this chromophore-containing peptide as CTPF throughout the manuscript. We recently discovered that, although 405 nm photoexposure renders the β-barrel non-fluorescent, the solution exhibited a faint red fluorescence under a 365 nm UV transilluminator, due to the photoreleased CTPF (**Figure 1A**).^18^ We exploit our discovery and propose a recombinant approach to render a protein/peptide fluorescent by transferring the chromophore from PhoCl1 to the target peptide or protein to be labelled (TPTL) (**Figure 1B**). We supplement our experimental studies with quantum mechanical/molecular mechanics (QM/MM) simulations. Finally, we employed the fluorescently labelled protein to investigate its liquid–liquid phase separation behavior, demonstrating the applicability of this labelling strategy for monitoring phase-separated states. To the best of our knowledge, this light activation-mediated generation of red fluorescence in a non-fluorescent protein through a transfer of post-translationally modified genetically encoded chromophore has not been reported elsewhere.

**Figure 1.**
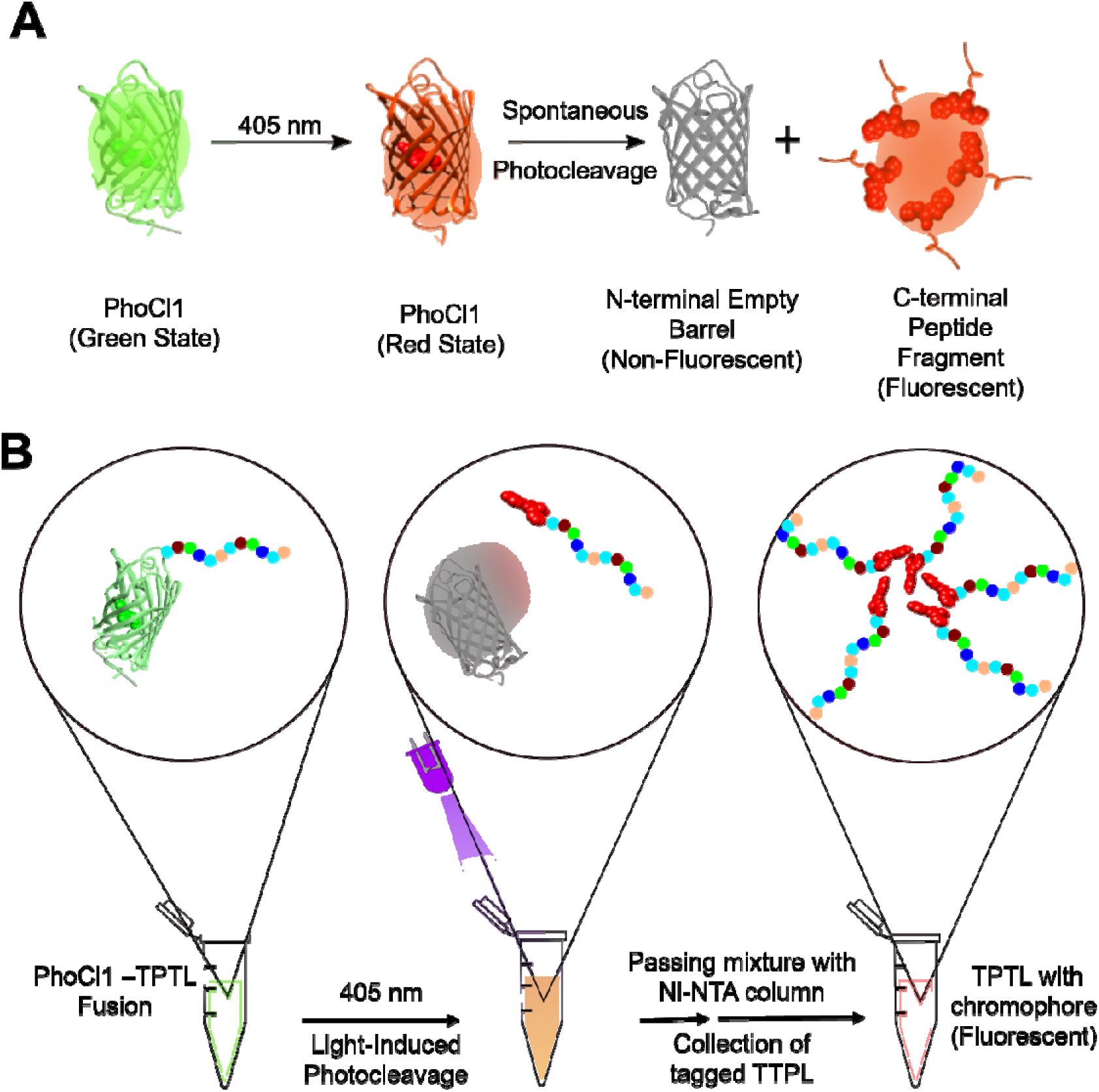
Photocleavage-based strategy using photo-cleavable protein (PhoCl1) enables fluorescent tagging of peptides/proteins after 405 nm light-induced dissociation. **(A)** PhoCl1 upon 405 nm photoexposure forms an intermediate in the red state, followed by its cleavage into 2 moieties: N-terminal empty barrel and C-terminal peptide fragment with the chromophore (CTPF). This CTPF is the source of red fluorescence, and **(B)** A fusion protein comprising PhoCl1 and the target peptide/protein to be labelled (TPTL) is expressed, and upon 405 nm photoexposure it will split into an N-terminal empty barrel and a fluorescent CTPF moiety fused to the TPTL. By using Ni-NTA chromatography, the 6xHis-tagged empty barrel will be retained on the resin, whereas the fluorescently labelled TPTL can be collected in the flow-through.

## RESULTS AND DISCUSSION

PhoCl1 undergoes photocleavage upon irradiation with 405 nm light, resulting in backbone cleavage yielding two distinct fragments: a non-fluorescent N-terminal empty β-barrel and a C-terminal peptide fragment that retains the chromophore (CTPF) (**Figure 1A**). We have previously shown that this CTPF exhibits red fluorescence.^18^ We propose to exploit this behavior for post-expression fluorescent labelling by genetically fusing PhoCl1 to the target peptide or protein to be labelled (TPTL) with a 6xHis tag in the N-terminal. Upon photoexposure, the fusion construct will undergo photocleavage resulting in a non-fluorescent, 6×His-tagged empty barrel and a fluorescent CTPF moiety covalently linked to the target peptide or protein. This will be subsequently purified using Ni–NTA affinity chromatography, which will selectively retain the His-tagged empty barrel on the resin, while the fluorescently labelled target peptide or protein will be recovered in the flow-through (**Figure 1B**). This strategy eliminates the need for excess small-molecule fluorophores and associated downstream purification steps, while also avoiding permanent fusion of large fluorescent proteins that can perturb native structure, function, or phase behavior of the labelled biomolecule.

We expressed two PhoCl1 variants with a 6×His tag positioned either at the N-terminus (6xHis-PhoCl1) or the C-terminus (PhoCl1-6xHis). These proteins were further immobilized onto the Ni–NTA resin. To validate light-induced photocleavage of PhoCl1 and the fluorescent properties of the released fragment, we compared the behaviour of these constructs before and after 405 nm irradiation (**Figure 2A**). The visual inspection of the supernatants revealed no detectable coloration for unexposed PhoCl1-6×His or 6×His-PhoCl1 samples. The photoexposed supernatant of the 6×His-PhoCl1 construct resulted in the appearance of a distinct red-colored supernatant corresponding to the released C-terminal peptide fragment (CTPF). In contrast, the empty barrel fragment remained colorless, consistent with its non-fluorescent nature in the photoexposed supernatant of the PhoCl1-6xHis. We ran the SDS-PAGE gel of photoexposed supernatants, i.e., CTPF and empty barrel, with controls, i.e., PhoCl1-6xHis and 6xHis-PhoCl1. The SDS–PAGE of CTPF (lane 2) showed no bands, suggesting there is no PhoCl1 or empty barrel contamination in purified CTPF supernatant (**Figure 2B**). The purity of the CTPF was also validated by MALDI-TOF analysis which revealed a dominant peak at m/z 1501.784 (M+H^+^), close to the theoretically calculated molecular weight of 1500.688 Da (**Figure S1**). Additionally, no peaks due to PhoCl1 empty barrel (MW: 28259.99 kDa) and intact PhoCl1(MW: 29779.72 kDa) were observable in the 20-30 kDa range, indicating minimal contamination (**Figure S2**). Fluorescence spectroscopy of the supernatants demonstrated that only the CTPF-containing fraction exhibited strong red fluorescence, with an emission maximum centered around ∼570 nm (**Figure 2C**). These results confirm that red fluorescence originates exclusively from the released CTPF following photocleavage and not from the intact protein or empty barrel fragment. Collectively, these observations establish that the C-terminal fragment can be selectively separated from the non-fluorescent empty barrel, providing a foundation for post-expression fluorescent labelling.

**Figure 2.**
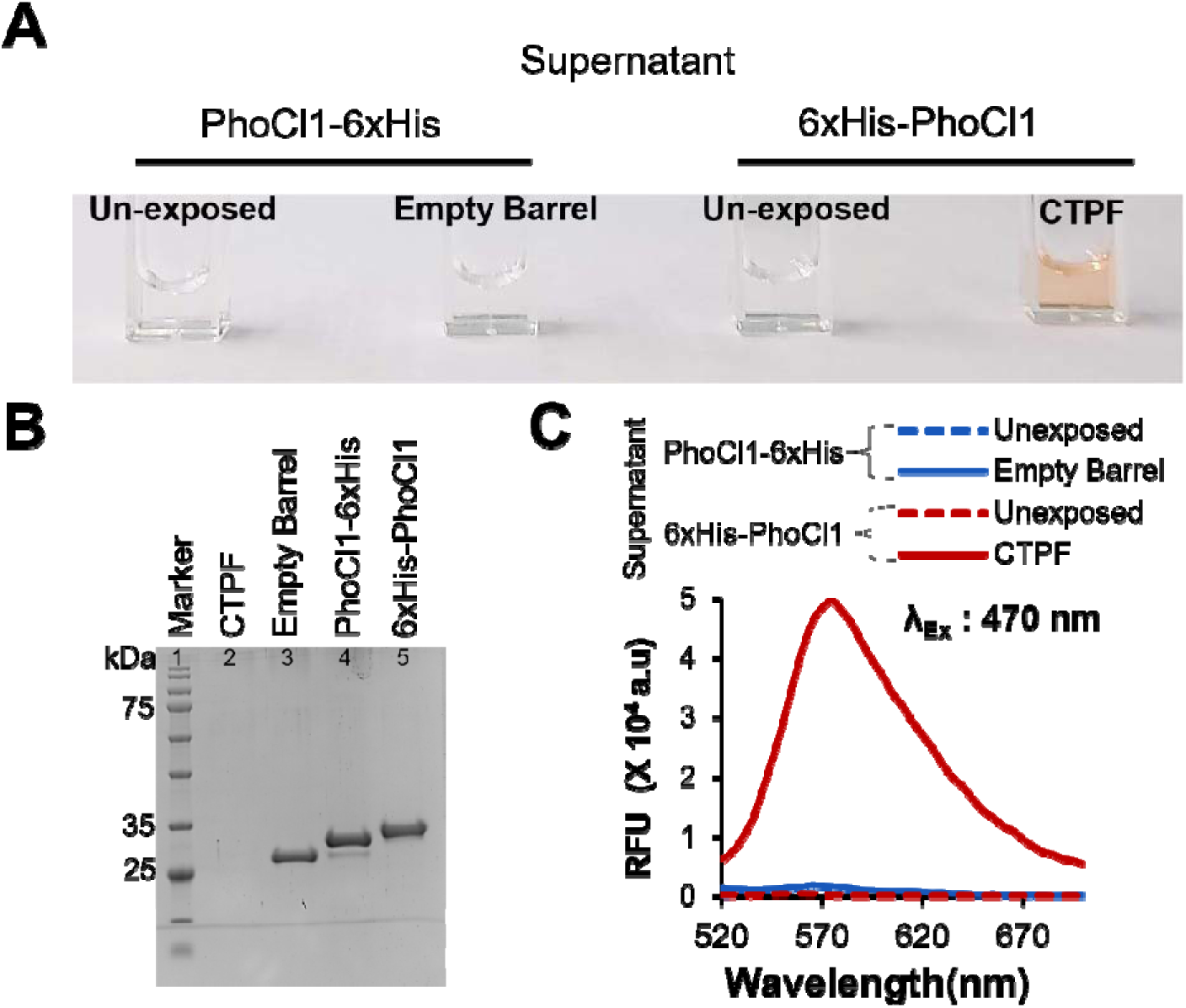
Photodissociation studies of 6xHis-PhoCl1 and PhoCl1-6xHis. **(A)** Visual image of the supernatants of PhoCl1-6xHis and 6xHis-PhoCl1 before and after photo-exposure, **(B)** 12% SDS-PAGE gel of supernatants of 6xHis-PhoCl1 (CTPF), PhoCl1-6xHis (PhoCl1 barrel) obtained after photo exposure, and controls PhoCl1-6xHis, and 6xHis-PhoCl1 and **(C)** Fluorescence emission spectra of PhoCl1-6xHis and 6xHis-PhoCl1 supernatants before and after photo exposure when excited at 470 nm.

Prior to tagging the TPTL with CTPF, we quantified the absorbance and fluorescence properties of CTPF under various conditions. CTPF absorbance spectra showed two major peaks, around 320 nm and 470 nm (**Figure S3**).^18^ To probe the influence of viscosity on CTPF fluorescence, we prepared a series of CTPF samples with increasing glycerol fractions (10%, 30%, 50%, and 70% v/v). An increase in fluorescence intensity was observed with increasing glycerol content, indicative of viscosity-dependent emission behavior (**Figure 3A and B**). This could be due to the restriction in intramolecular rotations within the chromophore at higher viscosities, thereby suppressing non-radiative decay pathways and enhancing radiative emission efficiency—an observation consistent with previously reported photophysical studies.^19, 20^ We also determined the fluorescence quantum yield (FQY) of CTPF in 70% glycerol solution, yielding a quantum yield of 1.91± 0.59 %, as compared to 0.57[±[0.07% of CTPF in the absence of glycerol.^18^

**Figure 3.**
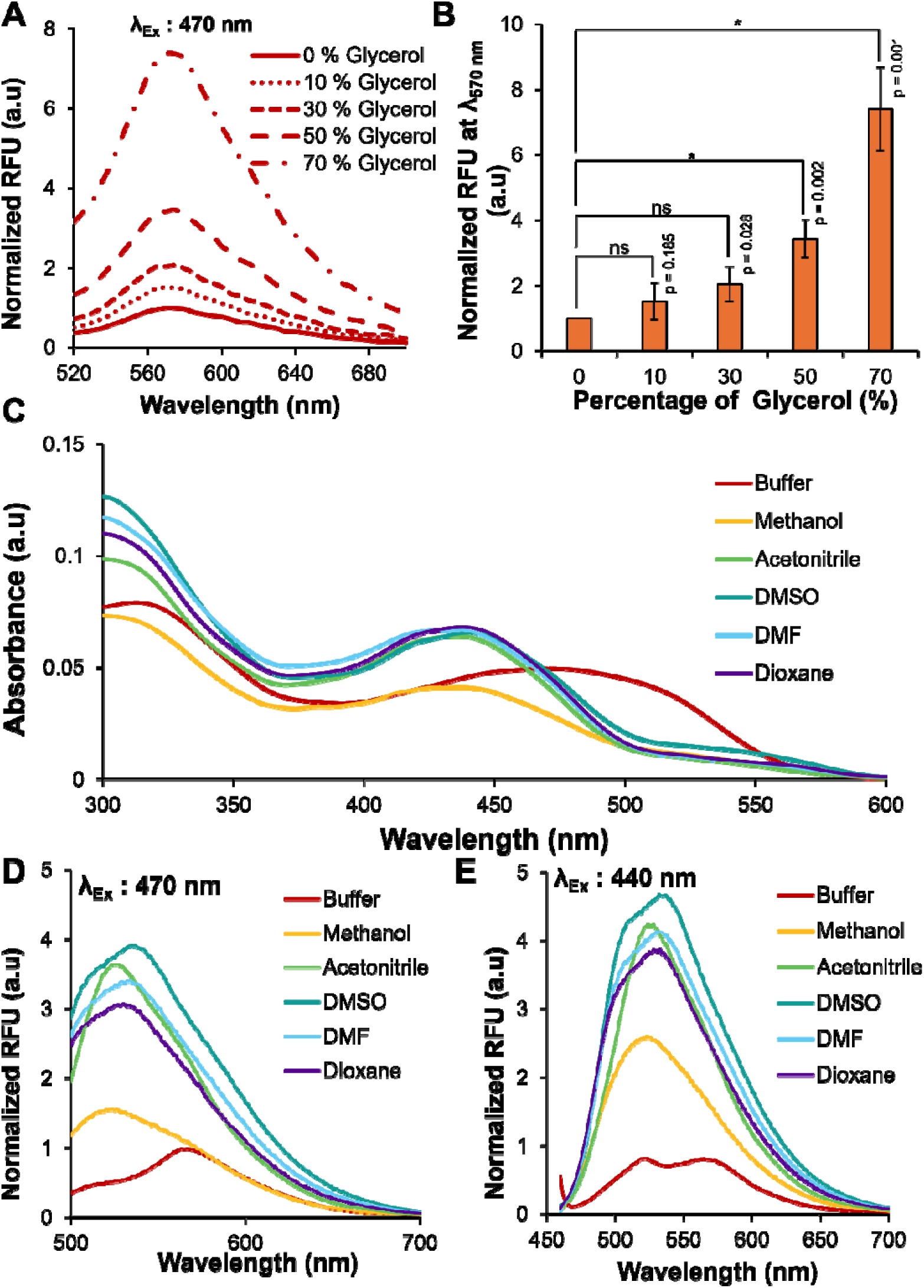
Spectral studies of CTPF under various solvents varying in viscosity and polarity. **(A)** Fluorescence emission spectra of CTPF in different concentrations of glycerol, **(B)** Peak emission intensity plotted as a function of increasing concentration of glycerol, **(C)** Absorbance spectra of CTPF in different solvents, **(D)** Fluorescence emission spectra of CTPF in different solvents when excited at 470 nm, and (**E**) Fluorescence emission spectra of CTPF in different solvents when excited at 440 nm.

Furthermore, we investigated the effect of increasing non-polarity on the fluorescence behavior of the CTPF aggregate. We recorded the absorbance and fluorescence spectra of CTPF in five solvents of increasing non-polarity^21^: Methanol (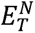 = 0.762) < Acetonitrile (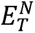 = 0.460) < Dimethyl Sulfoxide (DMSO) (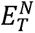 = 0.444) < Dimethylformamide (DMF) (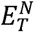 = 0.386) < Dioxane (£^N^ = 0.164). A distinct blue shift in the absorption maximum was observed, with the absorption maxima ∼470 nm in an aqueous buffer which shifted to ∼440 nm across all solvents examined (**Figure 3C**). Correspondingly, the fluorescence emission spectra exhibited both enhancement in the emission intensity and a hypsochromic shift, with the emission maximum decreasing from 570 nm in aqueous buffer to ∼540 nm in all the examined solvents (**Figures 3D and 3E**). Additionally, we investigated the effect on CTPF fluorescence in different surfactants. An increase in fluorescence emission intensity was observed with increasing detergent concentrations in all cases, accompanied by a blue shift, with the peak emission intensity shifting from 570 nm to ∼540 nm at higher concentrations of the detergent (**Figure S4**).

Having established the photophysical properties and release behavior of the C-terminal peptide fragment (CTPF), we next examined its utility as a fluorescent label for peptides and proteins. To evaluate the generality of this approach, we created fusion constructs of 6xHis-PhoCl1-TPTL. The expressed fusion was bound to Ni-NTA resin, photoexposed, and the released CTPF-TPTL was collected. This strategy was applied to a structurally and functionally diverse set of peptides and proteins.

As a validation model, we selected the silica-binding peptide R5 as the TPTL. R5 is a 19-residue peptide (SSKKSGSYSGSKGSKRRIL) derived from silaffin protein and known for its strong affinity toward silica, promoting both binding and precipitation.^22^ It has been widely explored in biotechnology for silicification-based encapsulation of enzymes,^23^ small molecules,^24^ and proteins, enabling applications in hybrid materials^25^ and biosensors.^22^ We created a fusion construct of PhoCl1 with R5 with a N-terminal 6xHis tag (6xHis-PhoCl1-R5). We followed the above-mentioned protocol of expression-binding-collection post photo-exposure, resulting in the collection of the red-colored supernatant consisting of CTPF-R5 (**Figure 4A**). We have performed MALDI-TOF analysis to validate the purity of the sample, which displayed an m/z peak at 4096.87, close to the theoretically calculated molecular weight of 4082.62 Da of CTPF-R5 (**Figure 4B and S5**). Additionally, no peaks due to PhoCl1 empty barrel (MW: 28259.99 kDa) in the 10-30 kDa range and the fusion protein, PhoCl1-R5 (MW: 31999.24 kDa) in the 30-50 kDa range were observed in MALDI-TOF, indicating minimal contamination (**Figure S6 and S7**). This was also corroborated with the SDS-PAGE analysis of purified CTPF-R5, which displayed an empty lane (difficulty in resolving small peptide fragments), confirming the absence of the fusion protein or the PhoCl1 empty barrel (**Figure 4C**). These taken together suggest that the red fluorescence is solely due to the CTPF-R5.

**Figure 4.**
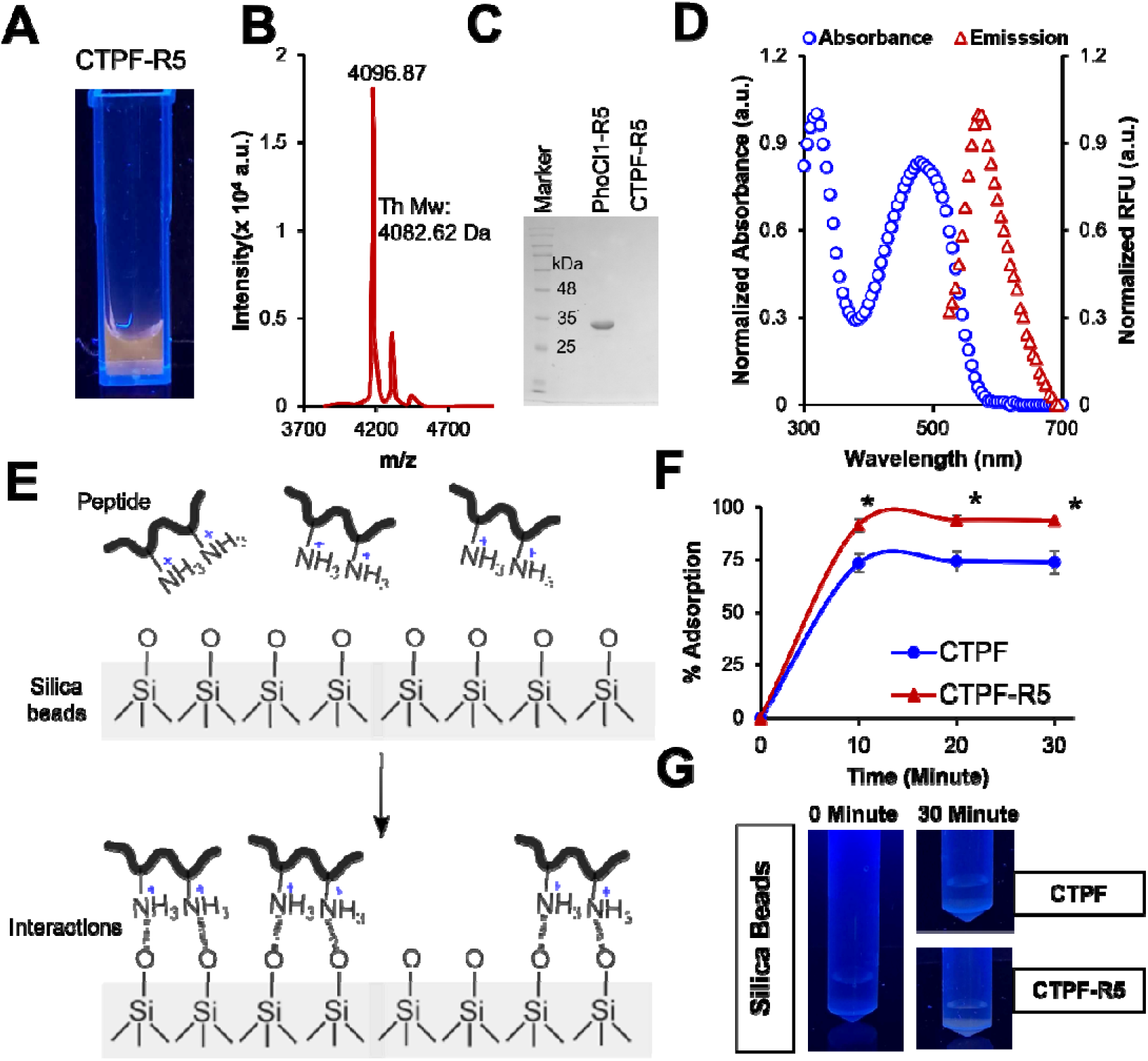
Conferring red fluorescence to non-fluorescent silica-binding peptide (R5) by photo-induced tagging with CTPF. **(A)** Visual image of purified CTPF-R5 acquired under a UV transilluminator (365 nm) displaying red-fluorescence, **(B)** Digitized MALDI-TOF mass spectrum of the purified CTPF-R5 showing a prominent peak at 4096.87 m/z, corresponding to the expected molecular weight (4082.62 Da) of CTPF-R5 (**Figure S5**), **(C)** 12% SDS-PAGE gel electrophoresis of unexposed PhoCl1-R5 and CTPF-R5. The purified R5 lane shows the absence of both PhoCl1-R5 and the PhoCl1 barrel, indicating the high purity of CTPF-R5, **(D)** Absorption spectra (blue circles) of the purified CTPF-R5 and emission spectra (red triangles) of the purified CTPF-R5 using excitation wavelength of 470 nm, **(E)** Schematic indicating the affinity of charged basic residues of the peptide binding to the negatively charged silica particles used to determine the activity of CTPF-R5 **(F)** Adsorption kinetics of CTPF and CTPF-R5, suggest adsorption of R5 on silica is significantly more than CTPF and (**G)** UV-transilluminator images of silica beads before adsorption (0 minute) and after adsorption (30 minute).

We have further characterized the CTPF-R5 peptide by acquiring absorbance and fluorescence spectra (**Figure 4D**). Upon excitation with 470 nm, the emission maximum was found to be ∼570 nm (**Figure 4D**). To assess whether the CTPF moiety affects the activity of the R5 domain, we performed silica-binding pull down assay based on electrostatic interactions between positively charged residues of R5 and the negatively charged silica surface (**Figure 4E**). Notably, CTPF-R5 exhibited significantly greater adsorption than CTPF alone (**Figure 4F**). This is further supported by the distinct red fluorescence observed on silica particles incubated with CTPF-R5 (**Figure 4G**). We corroborated the peptide binding strength by monitoring the red fluorescence in the supernatant following a buffer wash. CTPF was released in significantly higher amounts than CTPF-R5, indicating weaker binding (**Figure S8**). We further performed competitive elution using lysine, which disrupts the peptide interactions with silica by competing for binding.25 At 500 mM lysine, bound peptides were eluted, with CTPF showing greater release compared to CTPF-R5, confirming its weaker association with silica. A subsequent 2 M lysine wash resulted in complete desorption of both CTPF and CTPF-R5 peptide.

To expand the photo-tagging strategy to larger biomolecules, we generated a fusion protein with a riboflavin kinase as the fusion partner to PhoCl1 (6xHis-PhoCl1-*Mj*RibK). *Mj*RibK is an enzyme riboflavin kinase which was characterized from an archaea *Methanocaldococcus jannaschii*.26,27 It is a CTP-dependent enzyme involved in the biosynthesis of FMN (flavin mononucleotide) from riboflavin (vitamin B2).26,27 After photoexposure of 6xHis-PhoCl1-*Mj*RibK, the 6xHis-PhoCl1 empty barrel was bound to Ni-NTA beads while unbound CTPF-*Mj*RibK was collected. As anticipated, the expression-binding-collection post photo-exposure of 6xHis-PhoCl1-*Mj*RibK resulted in the generation of a red colored supernatant (**Figure 5A**). This was further assessed for its purity, molecular weight, and enzymatic activity. The 15 % SDS-PAGE gel electrophoresis of the supernatant showed a band at ∼17 kDa corresponding to the molecular weight of CTPF-*Mj*RibK (**Figure 5B**). The MALDI-TOF analysis also displayed an m/z peak at 17640.569 **(Figure 5C and Figure S9)** closely matching the theoretically predicted mass (17649.688 Da) of CTPF-*Mj*RibK with the absence of peaks corresponding to the intact fusion protein PhoCl1-*Mj*RibK and empty PhoCl1 barrel (**Figure S10)**. Next, we investigated the spectral characteristics of CTPF-*Mj*RibK through UV-Vis and fluorescence spectroscopy. We recorded the emission spectrum of CTPF-*Mj*RibK at the maximum excitation wavelength, which depicted an emission profile with a peak centered at ∼570 nm (**Figure 5D**).

**Figure 5.**
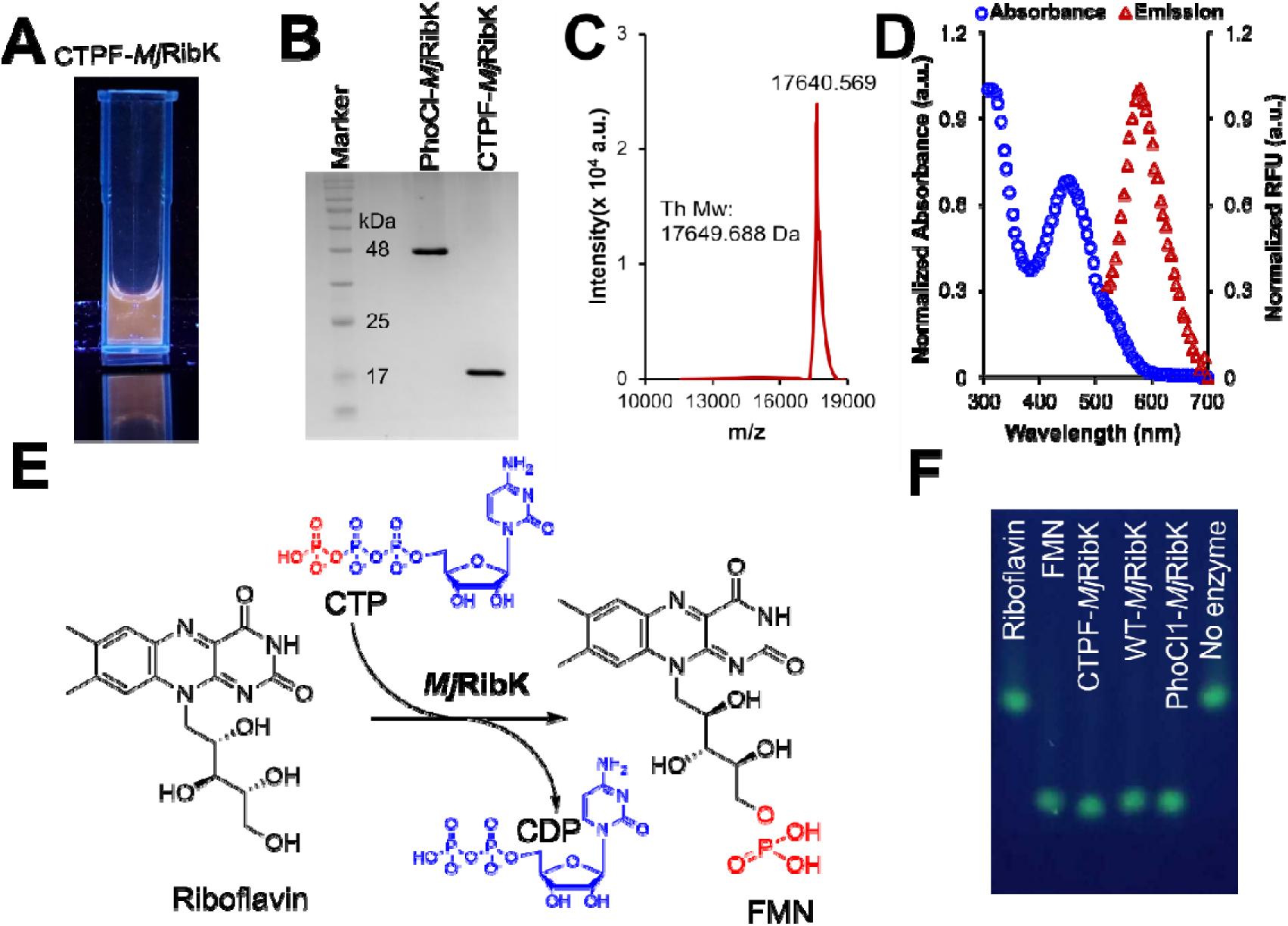
Conferring red fluorescence to non-fluorescent enzyme *Mj*RibK by photo-induced tagging with CTPF. **(A)**. Visual image of purified CTPF- *Mj*RibK acquired under a UV transilluminator (365 nm) displaying red-fluorescence, **(B)** 15 % SDS PAGE gel electrophoresis of unexposed PhoCl1-*Mj*RibK and CTPF- *Mj*RibK. The first lane is the molecular weight marker. The purified CTPF-*Mj*RibK lane shows the absence of both PhoCl1- *Mj*RibK and the PhoCl1 barrel. A single band at ∼17 kDa is present indicating the high purity of CTPF- *Mj*RibK, **(C)** Digitized MALDI-TOF mass spectrum of the purified CTPF-*Mj*RibK showing a prominent peak at 17640.569 m/z, corresponding to the theoretical molecular weight (17649.688 Da) of the peptide (**Figure S9**), **(D)** Absorption spectrum (blue circles) of the purified CTPF- *Mj*RibK and emission spectrum (red triangles) of the purified CTPF- *Mj*RibK using excitation wavelength of 470 nm, **(E)** Schematic representing reaction catalyzed by *Mj*RibK. It involves biosynthesis of FMN (flavin mononucleiotide) from riboflavin (vitamin B2) using CTP (Cytidine triphosphate) as a phosphate donar and releases CDP (Cytidine diphosphate) and **(F)** Fluorescent TLC image displaying the standards and riboflavin kinase activity of the fluorescently tagged enzyme. The first and second lanes correspond to riboflavin (RF) and flavin mononucleotide (FMN) standards respectively. The third and fourth lanes display a single fluorescence spot indicating the conversion of RF into FMN by CTPF-*Mj*RibK and WT-*Mj*RibK respectively. The fifth and sixth lanes are the PhoCl1-*Mj*RibK reaction and no enzyme controls.

We validated the retainment of enzymatic activity of CTPF-*Mj*RibK involving the phosphorylation of riboflavin into flavin mononucleotide (FMN) using cytidine-5’-triphosphate (CTP) as a phosphate donor (**Figure 5E**). By leveraging the distinct migration rates of riboflavin (Lane 1, retention factor = 0.5) and FMN (Lane 2, retention factor = 0.2), we successfully separated these compounds on a thin-layer chromatography (TLC) plate using the upper layer of solvent-mixture-n-butanol, acetic acid and water in 4:1:5 ratio (**Figure 5F**).28 The intrinsic fluorescence of riboflavin and FMN allowed us to visualize them on a TLC viewer instrument coupled with long range UV light (365 nm). Notably, the enzymatic reactions with CTPF-*Mj*RibK (Lane 3), WT-*Mj*RibK (Lane 4), and PhoCl1-*Mj*RibK fusion (Lane 5) displayed a single spot on the TLC plate at a retention factor of 0.2, indicating complete conversion of riboflavin to FMN. These revealed no variations in enzyme activity between them, while the no enzyme control (Lane 6) displayed no fluorescence intensity at the corresponding FMN spot. These taken together indicate that the activity of CTPF-*Mj*RibK was maintained and similar to WT-*Mj*RibK. Finally, we extended the photo-tagging approach to other peptides and proteins, including maltose binding protein (MBP), tobacco etch virus (TEV) protease, and a lanthanide-binding peptide (LBT). A distinct red fluorescence was observed under the UV transilluminator, with similar absorption and emission profiles. This was supported by SDS-PAGE and MALDI analyses, which confirmed the presence of a red peptide/protein (**Figure 6**).

**Figure 6.**
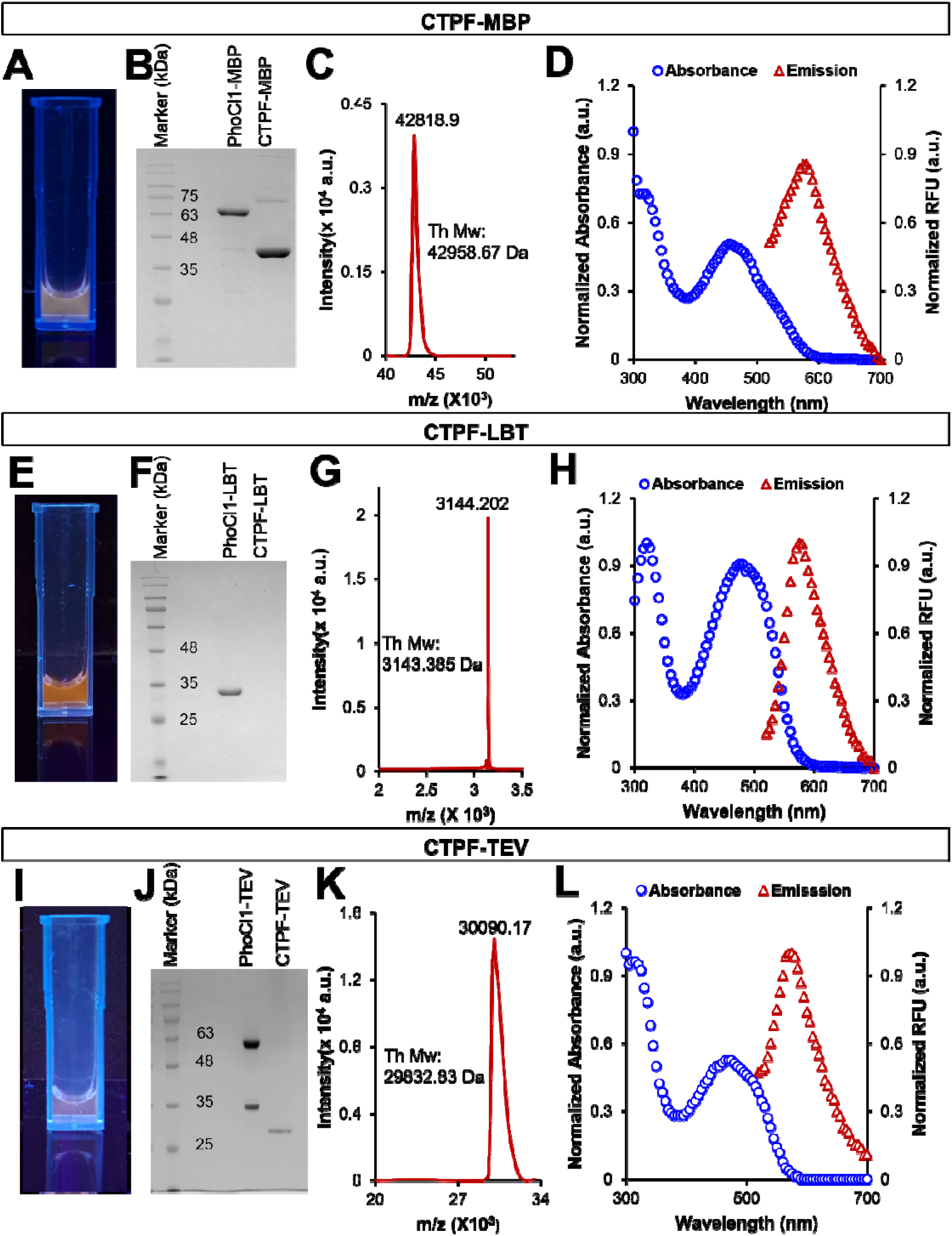
Conferring red fluorescence to non-fluorescent peptides and proteins by light-induced tagging with CTPF. **(A)** Visual image of purified CTPF-MBP acquired under a UV transilluminator (365 nm) displaying red-fluorescence, **(B)** 12% SDS-PAGE gel electrophoresis of unexposed PhoCl1-MBP and CTPF-MBP. The purified CTPF-MBP lane shows the presence of a band around 42 kDa corresponding to the theoretical molecular weight of CTPF-MBP of 42.958 kDa, **(C)** Digitized MALDI-TOF mass spectrum of the purified CTPF-MBP showing a prominent peak at 42818.9 m/z, corresponding to the theoretical molecular weight (42958.67 Da) of the protein (**Figure S11**), **(D)** Absorption spectrum (blue circles) and emission spectrum (red triangles) of the purified CTPF-MBP, **(E)** Visual image of purified CTPF-LBT acquired under a UV transilluminator (365 nm) displaying red-fluorescence, **(F)** 12% SDS-PAGE gel electrophoresis of unexposed PhoCl1-LBT and CTPF-LBT. The purified LBT lane shows the absence of both PhoCl1-LBT and the PhoCl1 barrel, indicating the purity of CTPF-LBT, **(G)** Digitized MALDI-TOF mass spectrum of the purified CTPF-LBT showing a prominent peak at 3144.202 m/z, corresponding to the theoretical molecular weight (3143.358 Da) of the peptide **(Figure S14)**, **(H)** Absorption spectrum (blue circles) and emission spectrum (red triangles) of the purified CTPF-LBT, **(I)** Visual image of purified CTPF-TEV acquired under a UV transilluminator (365 nm) displaying red-fluorescence, **(J)** 12% SDS-PAGE gel electrophoresis of unexposed PhoCl1-TEV and CTPF-TEV. The purified CTPF-TEV lane shows the presence of a band around 29 kDa corresponding to the theoretical molecular weight of CTPF-TEV of 29.83 kDa, **(K)** Digitized MALDI-TOF mass spectrum of the purified CTPF-TEV showing a prominent peak at 30090.168 m/z, corresponding to the theoretical molecular weight (29832.828 Da) of the protein **(Figure S16)** and **(L)** Absorption spectrum (blue circles) and emission spectrum (red triangles) of the purified CTPF-TEV. Note: MALDI data validating the absence of PhoCl1 empty barrel and fusion contamination in CTPF-MBP, CTPF-LBT, and CTPF-TEV are provided in the supplementary figures (**Figures S12, S13, S15, and S17**)

We previously demonstrated that dynamic aggregation of the photo-detached CTPF led to the exclusion of water molecules which facilitated the observation of red emission.^18^ To validate the same phenomenon in the case of CTPF-R5 we performed dynamic light scattering, which indicated a hydrodynamic size of 174 ± 52 nm (**Figure 7A**). A similar dynamic aggregation propensity of CTPF-R5 was observed in our molecular dynamics (MD) simulations spanning 600 ns (**Figure 7B and C**). The total interaction energy of CTPF-R5 was determined to be approximately **-**6800 kJ/mol (**Figure 7D**). The number of water molecules around the aggregated chromophore was also determined to lie between 8-20 compared to the isolated chromophore (30) (**Figure 7E**).

**Figure 7.**
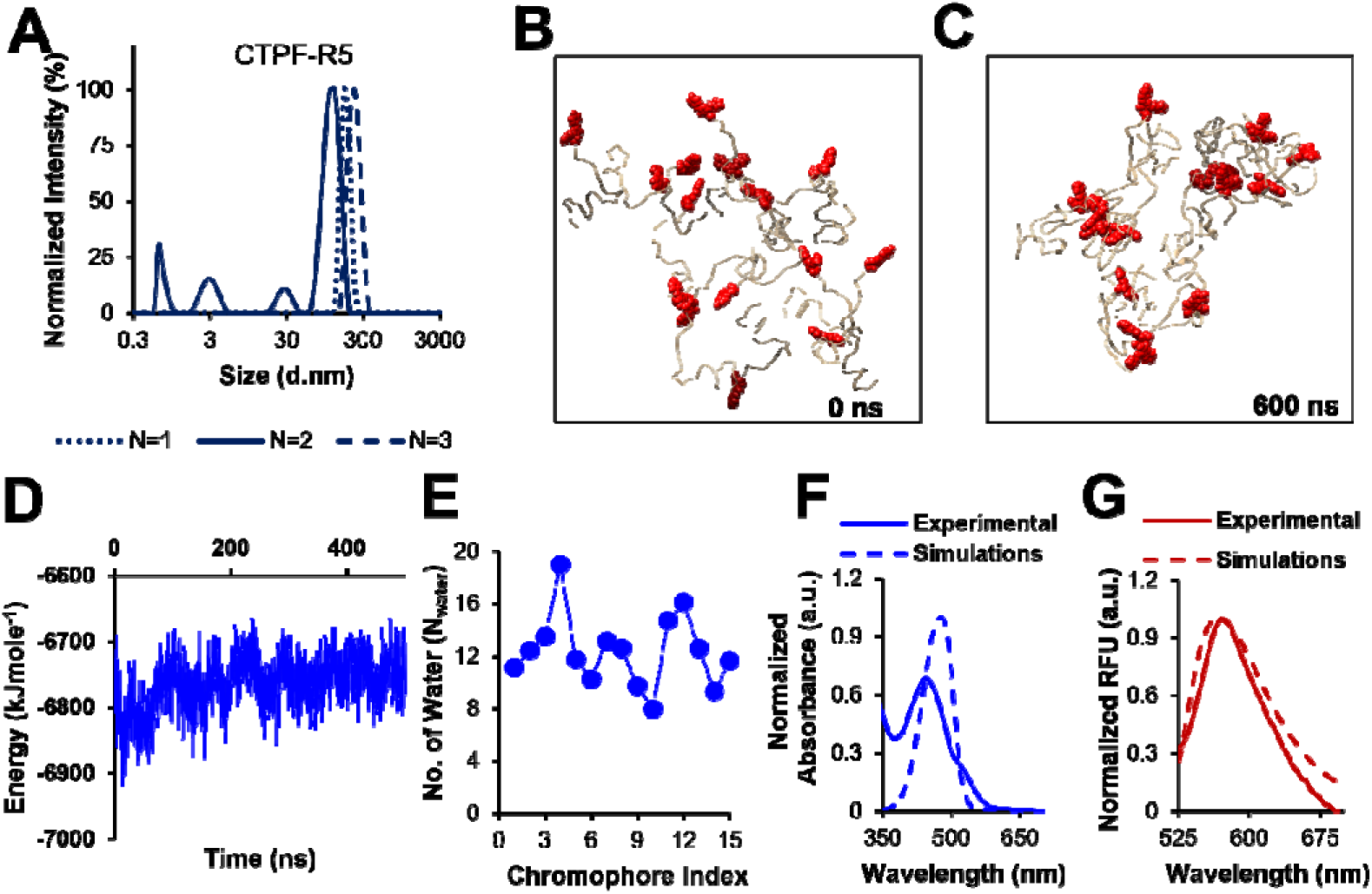
Size analysis of *in situ* generated CTPF-R5 and computational validation of red fluorescence observation supporting aggregation mechanism. **(A)** Size of CTPF-R5 was found to be 174 ± 52 nm, **(B)** Molecular dynamics snapshots of chromophore aggregation at time 0 ns, **(C)** Molecular dynamics snapshots of chromophore aggregation at time 600 ns. The chromophore is shown in red using a van der Waals representation, while the peptide is represented in golden, **(D)** Change in total interaction energy depending on time among all chromophores in molecular dynamics trajectory, **(E)** Number of water molecules (hydration number) around chromophores in 15 CTPF-R5 molecules, **(F)** Absorbance and **(G)** Emission scan acquired through QM/MM (dashed lines) by considering hydration number – overlayed with experimentally observed scans (solid lines) of CTPF-R5.

We performed quantum mechanics/molecular mechanics (QM/MM) calculations to get insight into the absorbance and fluorescence of the chromophore in its aggregated conformation. In the QM/MM setup, the chromophore was included in the QM region, while water molecules and residues within 3.5 Å of the chromophore were defined as the active region. The remaining residues were treated using the MM (molecular mechanics) approach. Initially, the entire system was minimized using quantum mechanical methods, after which the excitation and emission profiles were determined. As anticipated, the QM/MM calculations identified distinct absorption signatures in the 460 - 490 nm range, closely matching the experimentally observed peak at 470 nm (**Figure 7F)**. The emission maxima were computationally predicted to span 540–590 nm, which aligns with the experimental values centered near 570 nm (**Figure 7G**). The correlation between theoretical and experimental spectra further suggests that chromophore aggregation contributes to the red fluorescence of CTPF-R5. Similar analyses were performed for CTPF-*Mj*RibK with similar outcomes (**Figure S18**).

Finally, we demonstrate the potential application of the CTPF as a fluorescent label to visualize liquid liquid phase separation (LLPS). To demonstrate this principle, we constructed a PhoCl1 fusion with Elastin Like Polypeptide (ELP) with a N-terminal histidine tag (6xHis-PhoCl1-ELP). The same process followed earlier was used to purify the CTPF-ELP. A distinct red fluorescence was observed under the UV transilluminator (**Figure 8A**), The purity of CTPF-ELP is supported by SDS-PAGE as well as MALDI (**Figure 8B, 8C, S19 and S20**). CTPF-ELP demonstrated similar absorption and fluorescence spectra as observed earlier in other protein/peptide systems (**Figure 8D**). To visualize LLPS, the CTPF-ELP was incubated with PEG, a well-known crowding agent.29 The ELP condensates were observed under a fluorescence microscope at different excitation wavelengths (488 nm and 514 nm). The emission was acquired between 500-532 nm and 525-600 nm, respectively. Surprisingly, the CTPF-ELP was visualizable in both the green and red channel (**Figure 8E**). To validate this observation, we recorded the emission spectra and various excitation wavelengths and indeed there is a presence of a 510 nm peak at lower excitation wavelengths while the 570 nm emission peak is dominant at higher excitation wavelengths (**Figure S21**). While unsurprisingly, ELP alone without any fluorescent tag was not visualizable in the fluorescent channels (**Figure 8F**). The pure CTPF, when incubated with PEG alone, did not show the presence of coacervates, suggesting the role of ELP (**Figure S22A**). To rule out potential contamination from residual PhoCl1 fusion protein—prompted by fluorescence observed in the 500–532 nm emission window—we incubated PhoCl1 with PEG as a control. Fluorescence was detected in both the green and red channels; however, the signals did not spatially overlap, confirming that the observed emission in the CTPF-ELP samples likely does not arise from PhoCl1 contamination **(Figure S22B)**. As an additional experiment, we also demonstrate LLPS formation using CTPF–R5 **(Figure S23**). We further quantified the average particle diameters of the coacervates formed in the two systems. These exhibited an average particle diameter of 2.24 ± 0.9 μm (CTPF-ELP) and 0.93 ± 0.3 μm (CTPF-R5), respectively, with the corresponding size distributions shown in **Figure S24**.

**Figure 8.**
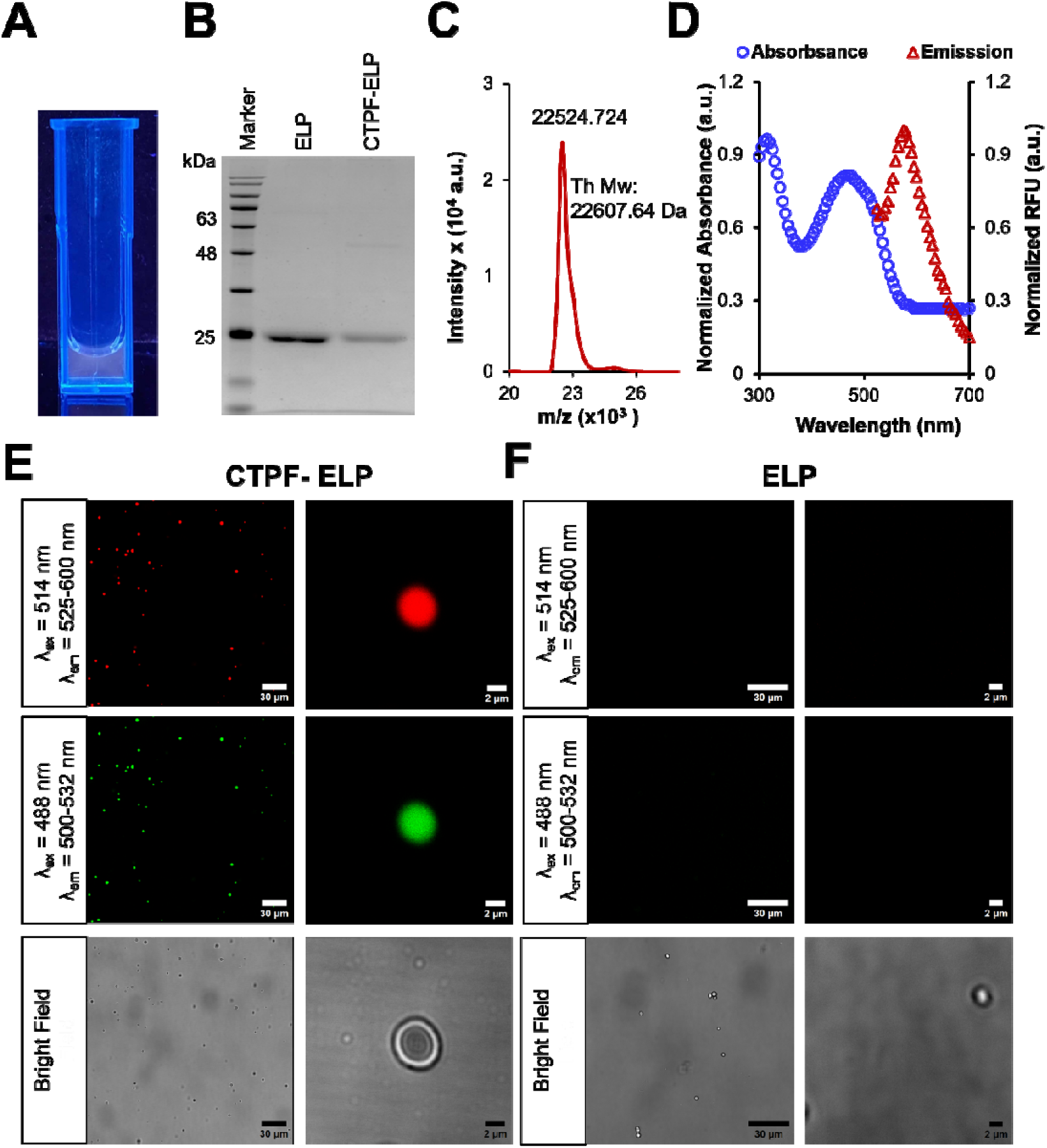
Tagging of Elastin Like Polypeptides with CTPF. **(A)** Visual image of purified CTPF-ELP acquired under a UV transilluminator (365 nm) displaying red-fluorescence, **(B)** 12% SDS-PAGE gel electrophoresis of purified ELP and CTPF-ELP, **(C)** Digitized MALDI-TOF mass spectrum of the purified CTPF-ELP showing a prominent peak at 22524.724 m/z, corresponding to the theoretical molecular weight (22607.638 kDa) of CTPF-ELP (**Figure S19**), **(D)** Absorption spectrum (blue circles) and emission spectrum (red triangles) of the purified CTPF-ELP, **(E)** Confocal images of CTPF-ELP observed under the red and green channels along with bright field micrographs and **(F)** Confocal images of ELP observed under the red and green channels along with bright field micrographs.

## CONCLUSIONS

In this study, we have demonstrated a methodology for fluorescent tagging of biomolecules (peptides and proteins) by leveraging the photocleavable property of PhoCl1. By recombinantly fusing PhoCl1 to the ‘target peptide/protein to be labeled’ (TPTL), we have achieved site-specific fluorescent labeling upon exposure to 405 nm light. Following purification, the labeled proteins and peptides were readily obtained without the need for additional purification steps. Functional assays confirmed that the labeling process does not significantly perturb the activity of the labeled peptides or proteins. We observed an increase in fluorescence intensity with increasing solution viscosity and a blue shift in the emission spectra upon varying solvent polarity. Quantum mechanics/molecular mechanics (QM/MM) calculations implicate the chromophore as a key contributor to the observed red emission. The generality of this approach was demonstrated across a broad range of peptides and proteins, including silica-binding and lanthanide-binding peptides, enzymes such as MjRibK and TEV protease, and proteins such as maltose-binding protein. Finally, we demonstrate the applicability of this recombinant photo-tagging strategy to elastin-like polypeptides, enabling direct visualization of phase separation without the need for small-molecule dyes or bulky fluorescent protein tags. We anticipate that this work will establish a simple, genetically encodable, and minimally perturbative photo-tagging platform that expands the toolkit for fluorescent labeling.

## MATERIALS AND METHODS

### DNA purification kits and enzymes

Plasmid purification kits, PCR purification kits, Gel extraction kits and Ni-NTA agarose were purchased from Qiagen. Q5® Hot Start High-fidelity 2X master mix, NEBuilder® HiFi DNA assembly master mix, T4 DNA Ligase, BamHI-HF®, NdeI, NcoI-HF®, KpnI-HF®, SacI-HF®, Dpn1, and XhoI restriction enzymes were acquired from New England Biolabs (NEB). PrimeSTAR GXL DNA polymerase was acquired from DSS Takara India Pvt. Ltd.

### Chemicals and reagents

Tris base, 2-mercaptoethanol, kanamycin sulfate, nickel sulfate, riboflavin, flavin mononucleotide (FMN), sodium chloride, n-butanol, acetic acid, methanol, glycerol, and bacterial growth media—Luria Bertani (LB) broth (Miller) and LB agar (Miller)—were obtained from Sisco Research Laboratory (SRL) Pvt. Ltd. Cytidine-5’-triphosphate (CTP) was sourced from Cayman Chemical (USA). Phenylmethylsulfonyl fluoride (PMSF) and disodium dihydrogen ethylenediaminetetraacetate dihydrate (EDTA) were supplied by Tokyo Chemical Industry (TCI) Co. Ltd. Isopropyl-ß-D-1-thiogalactopyranoside (IPTG), low EEO agarose, and imidazole were purchased from HiMedia Laboratories. Silica gel (Davisil Grade 643, pore size 150 Å, 200–425 mesh), and PEG (M_n_ 8000) was obtained from Merck.

### Consumables

Thin layer chromatography (TLC) plates (aluminum sheets coated with silica gel 60 F[[[) and 10 kDa MWCO Amicon® centrifugal filters were obtained from Merck. SnakeSkin™ dialysis tubing (10 kDa MWCO, 0.22 mm internal diameter) was procured from Thermo Fisher Scientific. UV LEDs (400–405 nm, model VAOL-5GUV0T4) were sourced from VCC Optoelectronics. Costar® 96-well UV-transparent microplates and black fluorescence microplates were purchased from Corning Incorporated.

### Plasmids

*Mj*RibK plasmid was a generous gift from Dr. Amrita Hazra (Whytamin Lab IISER Pune (India). *p*ET-PhoCl1-6xHis was a gift from Robert Campbell (Addgene plasmid #164033; http://n2t.net/addgene:164033; RRID: Addgene_164033). pET28-MBP-super TEV protease was a gift from Mark Howarth (Addgene plasmid # 171782 ; http://n2t.net/addgene:171782 ; RRID:Addgene_171782). V48 ELP was a gift from Andrew MacKay (Addgene plasmid # 68395 ; http://n2t.net/addgene:68395 ; RRID:Addgene_68395).

### Cloning of fusion constructs

The fusion of the PhoCl1 gene with the riboflavin kinase (*Mj*ribK) gene from *Methanocaldococcus jannaschii* in pET28a(+) vector was constructed using the NEB HiFi DNA assembly kit, following the manufacturer’s protocol. The fusion gene construct included a 6xHis tag sequence at the 5’ end of the PhoCl1 gene and the *Mj*RibK gene at the 3’ end, both connected via a flexible linker sequence KLGGGS. The NEB Q5®-Hot Start high-fidelity 2X master mix was used for PCR amplification, following the manufacturer’s instructions. We PCR amplified the gene sequences of PhoCl1 and *Mj*ribK with suitable overhangs using pET-PhoCl1-6xHis and pET28a-*Mj*RibK plasmids as templates respectively. Next, the pET28a(+) vector was linearized by digesting it with NcoI-HF and XhoI restriction enzymes. The linearized vector and the amplified gene sequences of PhoCl1 and *Mj*ribK were combined in a 1:2 ratio with the NEB HiFi DNA assembly master mix according to the company’s protocol and incubated at 50 °C for 15 min to facilitate DNA assembly. Subsequently, the DNA assembly was transformed into chemically competent *E. coli* DH5α cells. Plasmids were obtained from single colonies that grew after transformation.

The plasmid 6xHis-PhoCl1-R5 and 6xHis-PhoCl1-LBT has been constructed using restriction-free cloning.30 The 6xHis-PhoCl1 plasmid containing PhoCl1 gene was used as a template to PCR amplify the 6xHis-PhoCl1-R5 and 6xHis-PhoCl1-LBT gene sequence. Both fusion sequences were amplified such that it contained a flexible linker sequence (KLGGGS for R5 and GGGGS for LBT) between PhoCl1 and peptide gene. The PCR amplified fusion gene sequences were used as a megaprimer to conduct whole plasmid amplification which resulted in the insertion of the desired peptide sequence C-terminal to the PhoCl1 gene and separated by a flexible linker.

The amplified plasmid were treated with an enzyme DpnI to get rid of any parent template plasmid remaining. Finally, a digested product has been transformed into ultracompetent *E. coli* NEB DH5α cells.

The cloning of 6×His-PhoCl1-MBP, and 6×His-PhoCl1-TEV, was carried out in two main steps. First, restriction-free cloning was used to introduce compatible restriction sites at specific locations in both PhoCl1 and MBP-TEV plasmid constructs. (Few required restriction sites were already present in the construct of these and 6×His-PhoCl1V48ELP. 6×His-PhoCl1V48 ELP was transferred via digestion ligation to pET 28 a+, to introduce additional sites). For PhoCl1, a linker sequence (KLGGGS) flanked by KpnI and BamHI sites was incorporated by designing reverse primers containing the linker and restriction sites along with forward primers targeting the middle region of 6×His-PhoCl1. This generated a megaprimer that facilitated whole-plasmid amplification, resulting in a PhoCl1 construct with the linker and desired restriction sites. Similarly, KpnI and BamHI sites were introduced into the MBP-TEV plasmids around the MBP and TEV regions using restriction-free cloning. (While 6×His-PhoCl1 had the sites as it was in pET28 a+ plasmid). Next, the constructs were digested with appropriate restriction enzymes: MBP-TEV and PhoCl1-linker were digested with KpnI and SacI to prepare 6×His-PhoCl1-MBP, while BamHI and XhoI digestion was used to generate 6×His-PhoCl1-TEV. The digested MBP and TEV fragments were separated by gel electrophoresis and gel-extracted, whereas the PhoCl1-linker fragment was purified by PCR cleanup to retain the insert while removing small DNA fragments. The purified inserts were then ligated into the PhoCl1-linker vector at a 1:5 molar ratio using T4 DNA ligase. The ligation products were transformed into ultra-competent E. coli DH5α cells, and positive colonies were confirmed by plasmid purification, restriction digestion analysis, and sequencing (Barcode Bioscience). The resulting 6×His-PhoCl1-MBP, 6×His-PhoCl1-TEV and 6xHis-PhoCl1-ELP plasmids were subsequently used for protein expression.

Note: For restriction free cloning DNA Polymerase from DSS Takara was used.

### Overexpression of 6xHisPhoCl1, 6xHisPhoCl1-R5, 6xHis-PhoCl1-*Mj*RibK, 6xHis-PhoCl1-MBP, 6xHis-PhoCl1-TEV, 6xHis-PhoCl1-LBT and 6xHis-PhoCl1-ELP fusion protein

The plasmids 6xHisPhoCl1, 6xHisPhoCl1-R5, 6xHis-PhoCl1-*Mj*RibK, 6xHis-PhoCl1-MBP, 6xHis-PhoCl1-TEV, 6xHis-PhoCl1-LBT were transformed into *E. coli* BL21(DE3) chemical competent cells. The 1% of the overnight-grown transformed *E. coli* BL21(DE3) cultures of the above-mentioned plasmids were used to secondary inoculate the LB supplemented with 50 µg/mL kanamycin sulfate. This secondary culture was then incubated at 37 °C and 200 rpm in an incubator shaker. The optical density of the secondary culture was measured at 600 nm. Once it reached 0.6-0.7, induction was performed by adding 0.5 mM IPTG to promote overexpression. After induction, the temperature of the incubator was reduced to 18 °C and maintained for 18 h. These cells were harvested by centrifugation at 6,000 RPM at 4 °C for 10 min. The resulting cell pellets were flash-frozen and stored at - 80 °C for further use.

### Ni-NTA affinity purification of 6xHisPhoCl1, 6xHis-PhoCl1-R5, 6xHis-PhoCl1-*Mj*RibK Fusion, 6xHis-PhoCl1-MBP, 6xHis-PhoCl1-TEV, 6xHis-PhoCl1-LBT, and 6xHis-PhoCl1-ELP

The cell pellet was resuspended in lysis buffer (50 mM Tris-HCl pH 8, 300 mM NaCl, 0.1 mM PMSF). This was subjected to sonication using a cycle of 1 s ON, and 3 s OFF, 60 % amplitude, for total time of 20 minutes. Following sonication, the lysed cells were centrifuged at ∼16200 rpm for 50 minutes. The clarified lysate (supernatant) was collected and filtered with 0.22 micron syringe filter. Meanwhile, the Ni-NTA beads were equilibrated using the equilibration buffer (50 mM Tris-HCl pH 8, 300 mM NaCl). The filtered lysate was loaded onto the resin for binding and kept in refrigerator 4[C, 30 minutes. The column was washed with wash buffer (50 mM Tris-HCl pH 8, 300 mM NaCl, 20 mM imidazole). Finally, the protein was eluted in elution buffer (50 mM Tris-HCl pH 8, 300 mM NaCl, 150 mM imidazole). The eluted proteins were run on a 12% SDS gel to see the purity, and the pure proteins were dialyzed using a 10 kDa molecular weight cut off (MWCO) dialysis membrane in a buffer (50 mM Tris-HCl pH 8, 300 mM NaCl).

(Note: For 6xHis-PhoCl1-ELP, the Tris and NaCl content used was: 25mM Tris pH-8, 150 mM NaCl. Imidazole was the same as other constructs. And in 6xHis-PhoCl1-TEV, 10% Glycerol was present in the lysis, equilibration, and elution buffer, along with other components.)

### 405 nm light source configuration

A custom 405 nm light source was assembled using UV LEDs (400–405 nm, VAOL-5GUV0T4) to photoexpose the photocleavable protein PhoCl1 and its fusion constructs. The LED’s positive terminal was connected in series to a 470 Ω resistor, with both terminals mounted on a breadboard and powered by a 5 V DC supply. A schematic and full setup description is provided in our previous work.31 For photoexposure, the LED was positioned upside down on the breadboard and inserted into a microcentrifuge tube containing either PhoCl1, PhoCl1-R5, PhoCl1-*Mj*RibK, PhoCl1-MBP, PhoCl1-TEV, PhoCl1-LBT or PhoCl1-ELP.

### Isolation of pure CTPF, CTPF-R5, CTPF-*Mj*RibK, CTPF-MBP, CTPF-LBT, and CTPF-TEV and CTPF-ELP through 405 nm induced photocleavage

Purified 6xHis-tagged PhoCl1 was immobilized onto pre-equilibrated Ni-NTA affinity resin until saturation was achieved. The equilibration buffer consisted of 50 mM Tris-HCl (pH 8.0) and 300 mM NaCl. (Equilibration buffer used for ELP consisted of 25 mM Tris (pH 8.0) and 150 mM NaCl). After thorough washing to remove the unbound proteins, the resin with bound PhoCl1 was resuspended in fresh equilibration buffer and transferred into 2 mL microcentrifuge tubes (MCTs). After allowing the resin to settle (∼7.5 mm bead height), excess buffer was removed, leaving approximately 150 µL. The MCTs were then exposed to 405 nm light (1.5 mW·cm[²) for 12 hours using the custom LED setup. Post-irradiation, the supernatant containing the photo-released CTPF was collected. Similar procedure was used to get the other peptides or proteins including CTPF-R5, CTPF-*Mj*RibK, CTPF-MBP, CTPF-LBT, CTPF-TEV and CTPF-ELP.

Note: For CTPF-TEV, the temperature of 4□C was maintained surrounding MCT.

The purified CTPF, CTPF-R5, CTPF-*Mj*RibK, CTPF-MBP, CTPF-LBT, and CTPF-TEV were analyzed for molecular weight and purity using MALDI-TOF mass spectrometry. Additionally, 12% SDS-PAGE (15% gel for CTPF-*Mj*RibK) gel electrophoresis was performed to check for any higher molecular weight protein contaminants.

### Matrix-assisted laser desorption/ionization time-of-flight mass spectrometry (MALDI-TOF MS)

The purified protein/ peptide was buffer exchanged via washing fusion protein immobilized beads with 50 mM ammonium bicarbonate ((NH_4_)_2_CO_3_) buffer and following photoexposure the resulting supernatant containing POI was used for MALDI-TOF analysis. [For PhoCl1-TEV – the fusion protein immobilized beads were buffer exchanged with water, and the water was kept as supernatant also. After photoexposure, the supernatant of 2-3 tubes was pooled and lyophilized. The lyophilized samples were dissolved in 50 % acetonitrile and spotted on MALDI plate.]

A saturated solution of 2,5-dihydroxybenzoic acid (DHB) in a 30:70 mixture of acetonitrile and 0.1% trifluoroacetic acid (TFA) was prepared as the matrix. Equal volumes of matrix and protein samples were mixed and drop-casted on the MALDI target plate followed by drying at room temperature. The MALDI-TOF spectra were recorded on the Bruker Autoflex MALDI-TOF instrument in the m/z 1000-75000 for the protein CTPF-*Mj*RibK, CTPF-MBP, CTPF-TEV, CTPF-ELP, CTPF-R5, and CTPF-LBT. For the determination of the purity and molecular weight of CTPF-R5, α-cyano-4-hydroxycinnamic acid (CHCA) was used, and the spectra were recorded in the range m/z 1000-4500 in linear positive mode for CTPF-R5.

Note: For others except CTPF-R5, Linear positive mode has been used for m/z of 10000-75000, and reflectron mode has been used for m/z of 900 to 5000. The MALDI data were digitized using the online tool plotdigitizer.

### UV-Visible absorption and fluorescence emission spectra measurement

UV-Vis absorption and fluorescence emission spectra were recorded using a BioTek Cytation 5 multimode plate reader. For the absorption measurements, 100 µL of CTPF, CTPF-R5, CTPF-*Mj*RibK, CTPF-MBP, CTPF-LBT, CTPF-TEV and CTPF-ELP were placed separately in a 96-well UV-transparent microplate. Absorbance was measured across a wavelength range of 300 to 700 nm with a 5 nm step size. For fluorescence emission spectra, 100 µL of CTPF, CTPF-R5, CTPF-*Mj*RibK, CTPF-MBP, CTPF-LBT, and CTPF-TEV were loaded into a 96-well black polystyrene microplate. The excitation wavelength was kept 470 nm and emission was recorded over 520 to 700 nm.

### Transilluminator images capture

The transilluminator images of 90 µL purified peptides/ proteins were collected at 365 nm wavelength in a small plastic cuvette using a dual wave UV Transilluminator T4 from GeNei^TM^.

### UV-Visible Absorption and Fluorescence Emission Spectra Measurement for Solvatochromism and Viscosity Studies

The fluorescence emission spectra for viscosity studies were recorded using a BioTek Cytation 5 multimode plate reader. Fluorescence emission spectra of 150 μL sample containing, fixed CTPF (5 μM) concentration with increasing percentage of glycerol content (10%, 30%, 50%, and 70% v/v) was loaded into a 96-well black polystyrene microplate. The measurements were done using 470 nm excitation and emission from 520-700 nm. The emission spectra were normalized using the maximum emission intensity (λ = 570 nm) of the CTPF control. For the solvatochromism studies, 1000 μL of sample containing 70 % (v/v) different solvents individually with fixed CTPF concentration (5 μM) was used, the UV-Vis absorption spectra were measured from 300nm to 600 nm, at 5 mm spectral bandwidth, spectra were recorded using 1mm quartz cuvettes using a JASCO UV-Vis spectrophotometer (V-750). Identical samples composition was used to measure fluorescence emission studies using a Horiba-Jobin-Yvon Fluorolog-3 spectrofluorometer, spectra were recorded in quartz cuvettes with a 1 mm path length and slit widths of 5-7 nm. The measurements were done using two excitation wavelengths, 470 nm and 440 nm and emission from 490-700 nm and 460-700nm respectively. The emission spectra were normalized using the maximum emission intensity (λ = 570 nm) of the CTPF control.

### Quantum yield measurement of CTPF-R5 and CTPF-*Mj*RibK

UV-Vis absorption spectra were acquired using a JASCO V-750 spectrophotometer, while fluorescence emission measurements were conducted with a Horiba-Jobin-Yvon Fluorolog-3 spectrofluorometer. Spectra were recorded in quartz cuvettes with a 1 mm path length, using slit widths between 1 and 1.5 nm. Measurements were performed with the excitation wavelength matched to the sample’s absorption maximum. Fluorescein (quantum yield = 0.95 in 0.1 M NaOH) served as the reference standard for determining the quantum yield (QY) of CTPF. The QY was calculated using the following equation:

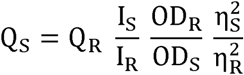

where Q is the quantum yield, I is the integrated fluorescence intensity, OD is the optical density, and η is the refractive index. Subscripts R and S denote the reference (fluorescein) and sample (peptide/ protein), respectively.

### Activity assays of CTPF-R5

#### Adsorption studies of CTPF-R5

A mixture of 20 μM, 200 μL of R5 peptide and CTPF as a supernatant was added to 24 mg of buffer equilibrated silica beads. The beads and peptide were mixed on a rotator. At each time point (10, 20, and 30 minutes), the supernatant was collected, and its fluorescence emission spectrum was recorded by exciting at 470 nm. After measurement, the supernatant was returned to the tubes, and the procedure repeated. The percentage adsorption at each time point was calculated from the initial fluorescence using the following equation:

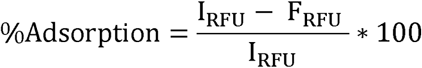

Where, I_RFU_ is the initial fluorescence intensity, F_RFU_ is the final fluorescence intensity at either of the time points 10, 20, or 30 minutes.

#### Assay of CTPF-R5

A mixture containing 20 μM R5 peptide and CTPF in 200 μL supernatant was added to 24 mg of buffer-equilibrated silica beads. After 30 minutes, the flow-through was collected. Subsequently, 200 μL of buffer (50 mM Tris-HCl, pH 8, 300 mM NaCl) was added, and the beads were rotated for another 30 minutes before collecting the supernatant. This process was repeated with buffers containing 500 mM lysine, followed by 2 M lysine, each with 30-minute incubation and collection steps.

Fluorescence emission measurements were performed on all collected supernatants.

#### Enzyme activity assay of WT-*Mj*RibK and CTPF-*Mj*RibK

The enzymatic activity of Red-*Mj*RibK was assessed through riboflavin kinase activity assay and compared with WT-*Mj*RibK.^1, 2^ To a mixture of 200 µM riboflavin, 50 mM Tris-HCl pH 8.0, 10 mM cytidine-5′-triphosphate (CTP), 1 mM MgCl_2_, 5 µM of CTPF-*Mj*RibK was added. The samples were incubated at 70 °C for 6 h and the conversion of riboflavin into FMN was estimated through thin-layer chromatography (TLC). We set up the same reaction using WT-*Mj*RibK as a control. 0.8 µL reaction mixtures along with the standards and controls were spotted on a TLC plate. The mobile phase used to run the TLC was the upper layer of the mixture of solvents, n-butanol, acetic acid, and water in a 4:1:5 ratio.^3^ The fluorescent spots on the TLC plate corresponding to the reactant ‘riboflavin’ and the product ‘FMN’ were captured on a TLC viewer equipped long UV light (il._365 nm_).

#### Dynamic light scattering measurements

Particle size analysis of CTPF-R5 and CTPF-*Mj*RibK were conducted using Malvern Zetasizer Dynamic Light Scattering (DLS) instrument.

#### Phase Separation of V48ELP, CTPF-ELP, and PhoCl1

A solution with PEG 30 % and proteins, approximately 9 μM, was mixed by pipetting and left for 24 hours at room temperature. 8 μL of the mixture is taken on a clean slide and covered with coverslips, and imaging was performed on all mixtures using a Leica DMI8 confocal microscope equipped with a 63×/1.4 NA oil-immersion objective. Laser used are 488 nm (4%), 514 nm (4%) for excitation. Coacervate sizes were measured using FiJi (ImageJ). Image was caliberated with scale bar and diameter was measured manually for individual coacervates.

#### Data analysis for experiments

All the experiments are performed in triplicates unless otherwise stated. The data analysis was carried out using Microsoft Excel. Student t-test (two-tailed) is performed wherever applicable (p<0.05 - significant). Theoretical molecular weight of the peptide or protein was calculated using the online Expasy ProtParam tool. Fiji (ImageJ) has been used for coacervate analysis. Data are represented as mean ± standard deviation. MS PowerPoint and Inkscape open source graphic editor were used for preparing and finalizing figures.

### Molecular dynamics simulations

#### Force field generation

The initial conformation of *Mj*RibK (PDB ID: 2VBU) and R5 sequence is created using the tleap module of Ambertools. The conformation of chromophore in the negatively charged state, along with the hydrogen, was generated from the PDB 7DNA.32 The restricted electrostatic potential (RESP) charges of chromophore were obtained via the RED module using the HF/6-311G* basis set.33 To simulate the peptide environment surrounding chromophore, the terminal of chromophore was capped with N-methyl. The total charge of chromophore was set to -1 and the charge on N-Methyl was set to 0 during charge calculations. Thereafter, the GAFF was used to generate the force field parameters of chromophore using the calculated RESP charges.34,35 Finally, the initial conformation and force field parameters for the entire conjugated system, including the chromophore, were generated using the XLeap module of AmberTools36. The AMBER formatted force field and coordinates were converted to the GROMACS formatted coordinates and topology using a Python script.37 All the simulation is performed using GROMACS formatted coordinates and topology using GROMACS-2020.2 version.38

#### Simulation Details

The conjugated system was placed in a cubic box of 120 Å dimensions. To eliminate any steric contact from the system, an energy minimization in vacuum was performed using the steepest descent method for 10000 steps with the tolerance force 0.001 kJmol^−1^. The system was solvated using the TIP3P water model, followed by the addition of 150 mM NaCl ions to neutralize the system.39,40 A second energy minimization was performed using the steepest descent method for 5,000 steps with a tolerance force of 0.001 kJmol^−1^. Thereafter, the system was heated to 27 °C using the Berendsen thermostat for 10 ns, during which the heavy atoms of the solute were position restrained.41 Subsequently, a 10 ns simulation was performed under constant temperature (27 °C) and pressure (1 atm) conditions, using a V-rescale thermostat and Berendsen barostat, respectively. This was followed by a 400 ns constant volume and temperature (NVT) production run at 300 K using the Nose–Hoover thermostat.41–44 During the simulation, the bonds were constrained using the LINC algorithm.45 The electrostatic effects were treated with the PME method using the 10 Å distance cut-off for long-range interaction.

Further, to investigate chromophore aggregation, total interaction energy calculations between all the chromophores were performed. The energy fluctuations converged, indicating the complete aggregation. The resulting aggregated structures were then analyzed using QM/MM simulations to determine the chromophore’s spectroscopic characteristics.

#### QM/MM Calculation

The spectroscopic properties of the chromophore were determined through QM/MM calculations on chromophore aggregates. The final aggregated conformation was selected based on the most stable energies obtained from molecular dynamics (MD) simulations. Subsequently, water molecules within a 3.5 Å radius of the chromophore were calculated to assess the hydration shell. Based on the hydration number, we have selected the chromophores for the QM/MM calculation.

In each QM/MM simulation (i) chromophore were treated in the quantum mechanical (QM) region (ii) residues within 3.5 Å of the chromophore comprised of the active region, including water (iii) remaining atoms were modeled using molecular mechanics (MM). The QM region is treated using B3LYP functional and cc-PVDZ basis set while the charges and van der Waals parameter of MM region is taken from the amber force field. The electrostatic at the QM//MM boundary is treated using the Electrostatic Embedding model.46,47 Initially, the aggregated conformation was optimized at the ground-state geometry level using the aforementioned parameters, followed by the calculation of the Hessian matrix to determine the vibrational levels of the system in its ground state. Subsequently, vertical excitation calculations were performed using Time-Dependent Density Functional Theory (TD-DFT) to identify the maximum excitation energy.48 Thereafter, the excited-state geometry was optimized and Hessian matrix at excited geometry was calculated to generate the excited-state vibrational energy levels. Vertical emission calculations were conducted to investigate fluorescence properties. All the QM/MM calculation are performed using the ORCA-5.0.0 package.49

#### Data Analysis

All experiments were carried out independently three times unless otherwise mentioned. Data analyses and plotting were carried out using Microsoft Excel and Origin plotting suit. Data are represented as mean ± standard deviation. All the MD simulation data was analyzed by using the various GROMACS modules and python script. The structural analysis of the proteins data bank (PDB) files was done using the visualization software’s ‘UCSF Chimera’ and ‘Visual molecular dynamics (VMD). Figures were prepared and finalized using Microsoft PowerPoint and Inkscape open-source vector graphic editor.

## Supporting information

https://drive.google.com/file/d/1SlnjAXvruzjoOacu2GvaGfv5Gtdx5rUy/view?usp=sharing

---

**Table of Contents Graphic.**
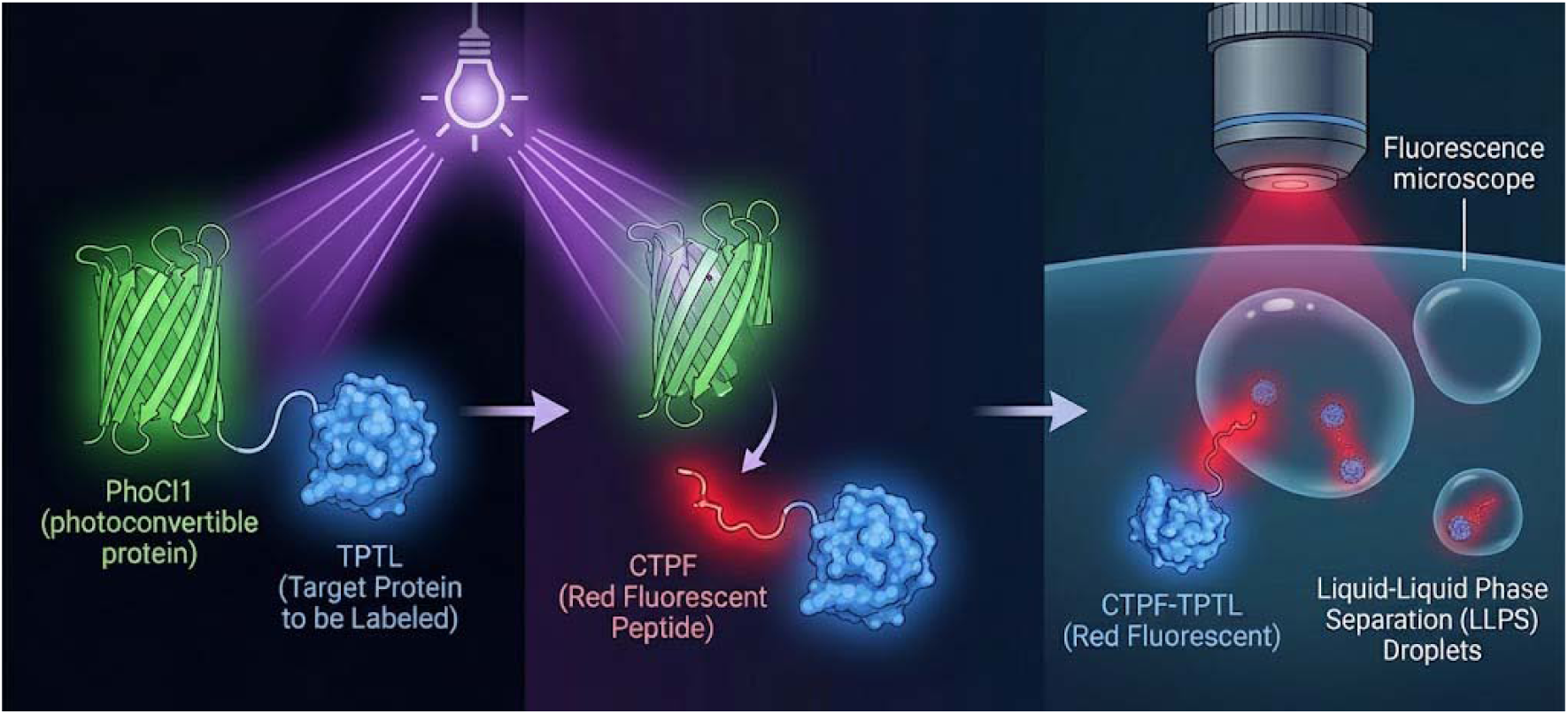
The TOC was generated through multiple prompts from various AI tools, and the final image was generated through Nano Banana Pro.

## AUTHOR CONTRIBUTIONS

KP conceived the original idea and planned the experiments. K.A, S.P, and Y.K. carried out the experiments and data analysis. R.K.S conceived, planned and analyzed the simulation data. K.P supervised the research along with providing feedback during the writing of the manuscript. All authors provided critical feedback and helped shape the research, analysis, and manuscript.

## ACKNOWLEDGEMENTS

The authors thank CRTDH (Common Resource and Technology Development Hub), CIF (Central Instrumentation Facility). The authors would also like to acknowledge Param Ananta Supercomputer Facility (IIT Gandhinagar) for providing the necessary computational facilities. We acknowledge the generous support of the Science and Engineering Research Board (SERB) and Startup Research Grant IIT Gandhinagar provided to K.P.

## Notes

### Competing Interest Statement

The authors have declared no competing interest.

