## Supplementary material for "Recombinant Photo-tagging Enables Fluorescent Labelling of Biomolecules and Visualization of Liquid–Liquid Phase Separation": https://drive.google.com/file/d/1SlnjAXvruzjoOacu2GvaGfv5Gtdx5rUy/view?usp=sharing

**Table S1.** Nucleotide/amino acid sequence of 6xHis-PhoCl1, 6xHis-PhoCl1-R5, 6xHis-PhoCl1-*Mj*RibK, 6xHis-PhoCl1-LBT, 6xHis-PhoCl1-MBP, 6xHis-PhoCl1-TEV, CTPF-R5, CTPF-*Mj*RibK, CTPF-LBT, CTPF-MBP and CTPF-TEV

**Text and other Color schemes:**

**Purple** - 6xHis tag  Dark Green - PhoCl1

**Grey Highlight (Bold)** -Chromophore forming residues

**Underline (Bold)** - Start codon and Stop codon

Red - Peptide / Protein of Interest

| **PhoCl1-6xHis nucleotide sequence** |
|---|
| **ATG**GTGATCCCTGACTACTTCAAGCAGAGCTTCCCCGAGGGCTACAGCTGGGAGCGC<br>AGCATGACCTACGAGGACGGCGGCATCTGCATCGCCACCAACGACATCACAATGGAG<br>GGGGACAGCTTCATCAACAAGATCCACTTCAAGGGCACGAACTTCCCCCCCAACGGCC<br>CCGTGATGCAGAAGAGGACCGTGGGCTGGGAGGCCAGCACCGAGAAGATGTACGAG<br>CGCGACGGCGTGCTGAAGGGCGACGTGAAGATGAAGCTGCTGCTGAAGGGCGGCGG<br>CCACTATCGCTGCGACTACCGCACCACCTACAAGGTCAAGCAGAAGCCCGTAAAGCTG<br>CCCGACTACCACTTCGTGGACCACCGCATCGAGATCCTGAGCCACGACAAGGACTACA<br>ACAAGGTGAAGCTGTACGAGCACGCCGTGGCCCGCAACTCCACCGACAGCATGGACG<br>AGCTGTACAAGGGTGGCAGCGGTGGCATGGTGAGCAAGGGCGAGGAGACCATTACAA<br>GCGTGATCAAGCCTGACATGAAGAACAAGCTGCGCATGGAGGGCAACGTGAACGGCC<br>ACGCCTTCGTGATCGAGGGCGAGGGCAGCGGCAAGCCCTTCGAGGGCATCCAGACGA<br>TTGATTTGGAGGTGAAGGAGGGCGCCCCGCTGCCCTTCGCCTACGACATCCTGACCA<br>CCGCCTTCCACTACGGCAACCGCGTGTTCACCAAGTACCCACGGGGAGGTGGAGGTC<br>TCGAGCACCACCACCACCACCAC**TGA** |
| **6xHis-Phocl1 amino acid sequence** |
| MVIPDYFKQSFPEGYSWERSMTYEDGGICIATNDITMEGDSFINKIHFKGTNFPPNGPVMQ<br>KRTVGWEASTEKMYERDGVLKGDVKMKLLLKGGGHYRCDYRTTYKVKQKPVKLPDYHFV<br>DHRIEILSHDKDYNKVKLYEHAVARNSTDSMDELYKGGSGGMVSKGEETITSVIKPDMKNK<br>LRMEGNVNGHAFVIEGEGSGKPFEGIQTIDLEVKEGAPLPFAYDILTTAF**HYG**NRVFTKYPR<br>GGGGLEHHHHHH |
| **6xHis-PhoCl1 nucleotide sequence** |
| **ATG**GGCAGCAGCCATCATCATCATCATCACAGCAGCGGCCTGGTGCCGCGCGGC<br>AGCCATATGGTGATCCCTGACTACTTCAAGCAGAGCTTCCCCGAGGGCTACAGCT<br>GGGAGCGCAGCATGACCTACGAGGACGGCGGCATCTGCATCGCCACCAACGACA<br>TCACAATGGAGGGGGACAGCTTCATCAACAAGATCCACTTCAAGGGCACGAACT<br>TCCCCCCCAACGGCCCCGTGATGCAGAAGAGGACCGTGGGCTGGGAGGCCAGCA<br>CCGAGAAGATGTACGAGCGCGACGGCGTGCTGAAGGGCGACGTGAAGATGAAG<br>CTGCTGCTGAAGGGCGGCGGCCACTATCGCTGCGACTACCGCACCACCTACAAG<br>GTCAAGCAGAAGCCCGTAAAGCTGCCCGACTACCACTTCGTGGACCACCGCATC<br>GAGATCCTGAGCCACGACAAGGACTACAACAAGGTGAAGCTGTACGAGCACGCC<br>GTGGCCCGCAACTCCACCGACAGCATGGACGAGCTGTACAAGGGTGGCAGCGGT<br>GGCATGGTGAGCAAGGGCGAGGAGACCATTACAAGCGTGATCAAGCCTGACATG<br>AAGAACAAGCTGCGCATGGAGGGCAACGTGAACGGCCACGCCTTCGTGATCGAG<br>GGCGAGGGCAGCGGCAAGCCCTTCGAGGGCATCCAGACGATTGATTTGGAGGTG<br>AAGGAGGGCGCCCCGCTGCCCTTCGCCTACGACATCCTGACCACCGCCTTCCACT<br>ACGGCAACCGCGTGTTCACCAAGTACCCACGG**TAA** |
| **6xHis-Phocl1 amino acid sequence** |
| MGSSHHHHHHSSGLVPRGSHMVIPDYFKQSFPEGYSWERSMTYEDGGICIATNDITM<br>EGDSFINKIHFKGTNFPPNGPVMQKRTVGWEASTEKMYERDGVLKGDVKMKLLLK |

GGGHYRCDYRTTYKVKQKPVKLPDYHFVDHRIEILSHDKDYNKVKLYEHAVARNS
TDSMDELYKGGSGGMVSKGEETITSVIKPDMKNKLRMEGNVNGHAFVIEGEGSGKP
FEGIQTIDLEVKEGAPLPFAYDILTTAFHYGNRVFTKYPR

**6xHis-PhoCl1-R5 nucleotide sequence**

**<u>ATG</u>**GGCAGCAGCCATCATCATCATCATCACAGCAGCGGCCTGGTGCCGCGCGGC
AGCCATATGGTGATCCCTGACTACTTCAAGCAGAGCTTCCCCGAGGGCTACAGCT
GGGAGCGCAGCATGACCTACGAGGACGGCGGCATCTGCATCGCCACCAACGACA
TCACAATGGAGGGGGACAGCTTCATCAACAAGATCCACTTCAAGGGCACGAACT
TCCCCCCCAACGGCCCCGTGATGCAGAAGAGGACCGTGGGCTGGGAGGCCAGCA
CCGAGAAGATGTACGAGCGCGACGGCGTGCTGAAGGGCGACGTGAAGATGAAG
CTGCTGCTGAAGGGCGGCGGCCACTATCGCTGCGACTACCGCACCACCTACAAG
GTCAAGCAGAAGCCCGTAAAGCTGCCCGACTACCACTTCGTGGACCACCGCATC
GAGATCCTGAGCCACGACAAGGACTACAACAAGGTGAAGCTGTACGAGCACGCC
GTGGCCCGCAACTCCACCGACAGCATGGACGAGCTGTACAAGGGTGGCAGCGGT
GGCATGGTGAGCAAGGGCGAGGAGACCATTACAAGCGTGATCAAGCCTGACATG
AAGAACAAGCTGCGCATGGAGGGCAACGTGAACGGCCACGCCTTCGTGATCGAG
GGCGAGGGCAGCGGCAAGCCCTTCGAGGGCATCCAGACGATTGATTTGGAGGTG
AAGGAGGGCGCCCCGCTGCCCTTCGCCTACGACATCCTGACCACCGCCTTCCACT
ACGGCAACCGCGTGTTCACCAAGTACCCACGGAAGCTTGGCGGCGGCTCTTCTTC
CTCTAAAAAGTCTGGTTCCTACTCTGGTAGCAAAGGCTCCAAACGTCGCATCCTG
**<u>TAA</u>**

**6xHis-PhoCl1-R5 amino acid sequence**

MGSSHHHHHHSSGLVPRGSHMVIPDYFKQSFPEGYSWERSMTYEDGGICIATNDITM
EGDSFINKIHFKGTNFPPNGPVMQKRTVGWEASTEKMYERDGVLKGDVKMKLLLK
GGGHYRCDYRTTYKVKQKPVKLPDYHFVDHRIEILSHDKDYNKVKLYEHAVARNS
TDSMDELYKGGSGGMVSKGEETITSVIKPDMKNKLRMEGNVNGHAFVIEGEGSGKP
FEGIQTIDLEVKEGAPLPFAYDILTTAFHYGNRVFTKYPRKLGGGSSSSKKSGSYSGS
KGSKRRIL

**6xHis-PhoCl1-*Mj*RibK nucleotide sequence**

**<u>ATG</u>**GGCAGCAGCCATCATCATCATCATCACGGCGGTACCGTGATCCCTGACTACT
TCAAGCAGAGCTTCCCCGAGGGCTACAGCTGGGAGCGCAGCATGACCTACGAGG
ACGGCGGCATCTGCATCGCCACCAACGACATCACAATGGAGGGGGACAGCTTCA
TCAACAAGATCCACTTCAAGGGCACGAACTTCCCCCCCAACGGCCCCGTGATGC
AGAAGAGGACCGTGGGCTGGGAGGCCAGCACCGAGAAGATGTACGAGCGCGAC
GGCGTGCTGAAGGGCGACGTGAAGATGAAGCTGCTGCTGAAGGGCGGCGGCCA
CTATCGCTGCGACTACCGCACCACCTACAAGGTCAAGCAGAAGCCCGTAAAGCT
GCCCGACTACCACTTCGTGGACCACCGCATCGAGATCCTGAGCCACGACAAGGA
CTACAACAAGGTGAAGCTGTACGAGCACGCCGTGGCCCGCAACTCCACCGACAG
CATGGACGAGCTGTACAAGGGTGGCAGCGGTGGCATGGTGAGCAAGGGCGAGG
AGACCATTACAAGCGTGATCAAGCCTGACATGAAGAACAAGCTGCGCATGGAGG
GCAACGTGAACGGCCACGCCTTCGTGATCGAGGGCGAGGGCAGCGGCAAGCCCT
TCGAGGGCATCCAGACGATTGATTTGGAGGTGAAGGAGGGCGCCCCGCTGCCCT
TCGCCTACGACATCCTGACCACCGCCTTCCACTACGGCAACCGCGTGTTCACCAA
GTACCCACGGAAGCTTGGCGGCGGCTCTTTGGTGAAATTGATGATTATTGAGGG
AGAAGTAGTTTCAGGACTTGGAGAAGGGAGATATTTTTTATCCCTCCCTCCTTAC
AAAGAGATATTTAAGAAGATTCTTGGCTTTGAACCTTATGAGGGGACATTAAATT
TAAAATTAGATAGAGAATTTGATATAAACAAATTTAAATATATTGAAACAGAGG
ATTTTGAATTTAATGGGAAAAGATTTTTTGGAGTTAAGGTTTTACCAATAAAAAT

| |
|---|
| ATTAATAGGTAATAAAAAAATAGATGGGGCGATAGTTGTGCCGAAAAAAACATA TCATAGTAGTGAGATTATAGAGATAATTGCCCCAATGAAACTTAGGGAGCAATT TAATTTAAAGGATGGAGATGTTATAAAAATACTAATTAAGGGAGATAAAGATGA A**TAA** |
| **6xHis-PhoCl1-*Mj*RibK amino acid sequence** |
| MGSSHHHHHHGGTVIPDYFKQSFPEGYSWERSMTYEDGGICIATNDITMEGDSFINKI HFKGTNFPPNGPVMQKRTVGWEASTEKMYERDGVLKGDVKMKLLLKGGGHYRCD YRTTYKVKQKPVKLPDYHFVDHRIEILSHDKDYNKVKLYEHAVARNSTDSMDELY KGGSGGMVSKGEETITSVIKPDMKNKLRMEGNVNGHAFVIEGEGSGKPFEGIQTIDL EVKEGAPLPFAYDILTTAF**HYG**NRVFTKYPRKLGGGSLVKLMIIEGEVVSGLGEGRY FLSLPPYKEIFKKILGFEPYEGTLNLKLDREFDINKFKYIETEDFEFNGKRFFGVKVLPI KILIGNKKIDGAIVVPKKTYHSSEIIEIIAPMKLREQFNLKDGDVIKILIKGDKDE |
| **6xHis-PhoCl1-LBT nucleotide sequence** |
| **ATG**GGCAGCAGCCATCATCATCATCATCACAGCAGCGGCCTGGTGCCGCGCGGC AGCCATATGGTGATCCCTGACTACTTCAAGCAGAGCTTCCCCGAGGGCTACAGCT GGGAGCGCAGCATGACCTACGAGGACGGCGGCATCTGCATCGCCACCAACGACA TCACAATGGAGGGGGACAGCTTCATCAACAAGATCCACTTCAAGGGCACGAACT TCCCCCCCAACGGCCCCGTGATGCAGAAGAGGACCGTGGGCTGGGAGGCCAGCA CCGAGAAGATGTACGAGCGCGACGGCGTGCTGAAGGGCGACGTGAAGATGAAG CTGCTGCTGAAGGGCGGCGGCCACTATCGCTGCGACTACCGCACCACCTACAAG GTCAAGCAGAAGCCCGTAAAGCTGCCCGACTACCACTTCGTGGACCACCGCATC GAGATCCTGAGCCACGACAAGGACTACAACAAGGTGAAGCTGTACGAGCACGCC GTGGCCCGCAACTCCACCGACAGCATGGACGAGCTGTACAAGGGTGGCAGCGGT GGCATGGTGAGCAAGGGCGAGGAGACCATTACAAGCGTGATCAAGCCTGACATG AAGAACAAGCTGCGCATGGAGGGCAACGTGAACGGCCACGCCTTCGTGATCGAG GGCGAGGGCAGCGGCAAGCCCTTCGAGGGCATCCAGACGATTGATTTGGAGGTG AAGGAGGGCGCCCCGCTGCCCTTCGCCTACGACATCCTGACCACCGCCTTCCACT ACGGCAACCGCGTGTTCACCAAGTACCCACGGGGCGGCGGCGGCTCCGATCCGG ATAAAGATGGCACCATTGATCTGAAAGAA**TAA** |
| **6xHis-PhoCl1-LBT amino acid sequence** |
| MGSSHHHHHHSSGLVPRGSHMVIPDYFKQSFPEGYSWERSMTYEDGGICIATNDITM EGDSFINKIHFKGTNFPPNGPVMQKRTVGWEASTEKMYERDGVLKGDVKMKLLLK GGGHYRCDYRTTYKVKQKPVKLPDYHFVDHRIEILSHDKDYNKVKLYEHAVARNS TDSMDELYKGGSGGMVSKGEETITSVIKPDMKNKLRMEGNVNGHAFVIEGEGSGKP FEGIQTIDLEVKEGAPLPFAYDILTTAFHYGNRVFTKYPRGGGGSDPDKDGTIDLKE |
| **6xHis-PhoCl1-MBP nucleotide sequence** |
| **ATG**GGCAGCAGCCATCATCATCATCATCACAGCAGCGGCCTGGTGCCGCGCGGC AGCCATATGGTGATCCCTGACTACTTCAAGCAGAGCTTCCCCGAGGGCTACAGCT GGGAGCGCAGCATGACCTACGAGGACGGCGGCATCTGCATCGCCACCAACGACA TCACAATGGAGGGGGACAGCTTCATCAACAAGATCCACTTCAAGGGCACGAACT TCCCCCCCAACGGCCCCGTGATGCAGAAGAGGACCGTGGGCTGGGAGGCCAGCA CCGAGAAGATGTACGAGCGCGACGGCGTGCTGAAGGGCGACGTGAAGATGAAG CTGCTGCTGAAGGGCGGCGGCCACTATCGCTGCGACTACCGCACCACCTACAAG GTCAAGCAGAAGCCCGTAAAGCTGCCCGACTACCACTTCGTGGACCACCGCATC GAGATCCTGAGCCACGACAAGGACTACAACAAGGTGAAGCTGTACGAGCACGCC GTGGCCCGCAACTCCACCGACAGCATGGACGAGCTGTACAAGGGTGGCAGCGGT GGCATGGTGAGCAAGGGCGAGGAGACCATTACAAGCGTGATCAAGCCTGACATG AAGAACAAGCTGCGCATGGAGGGCAACGTGAACGGCCACGCCTTCGTGATCGAG GGCGAGGGCAGCGGCAAGCCCTTCGAGGGCATCCAGACGATTGATTTGGAGGTG |

AAGGAGGGCGCCCCGCTGCCCTTCGCCTACGACATCCTGACCACCGCCTTCCACT
ACGGCAACCGCGTGTTCACCAAGTACCCACGGAAGCTTGGCGGCGGCTCTGGTA
CCATGGGAATCGAAGAAGGTAAACTGGTAATCTGGATTAACGGCGATAAAGGCT
ATAACGGTCTCGCTGAAGTCGGTAAGAAATTCGAGAAAGATACCGGAATTAAAG
TCACCGTTGAGCATCCGGATAAACTGGAAGAGAAATTCCCACAGGTTGCGGCAA
CTGGCGATGGCCCTGACATTATCTTCTGGGCACACGACCGCTTTGGTGGCTACGC
TCAATCTGGCCTGTTGGCTGAAATCACCCCGGACAAAGCGTTCCAGGACAAGCT
GTATCCGTTTACCTGGGATGCCGTACGTTACAACGGCAAGCTGATTGCTTACCCG
ATCGCTGTTGAAGCGTTATCGCTGATTTATAACAAAGATCTGCTGCCGAACCCGC
CAAAAACCTGGGAAGAGATCCCGGCGCTGGATAAAGAACTGAAAGCGAAAGGT
AAGAGCGCGCTGATGTTCAACCTGCAAGAACCGTACTTCACCTGGCCGCTGATTG
CTGCTGACGGGGGTTATGCGTTCAAGTATGAAAACGGCAAGTACGACATTAAAG
ACGTGGGCGTGGATAACGCTGGCGCGAAAGCGGGTCTGACCTTCCTGGTTGACC
TGATTAAAAACAAACACATGAATGCAGACACCGATTACTCCATCGCAGAAGCTG
CCTTTAATAAAGGCGAAACAGCGATGACCATCAACGGCCCGTGGGCATGGTCCA
ACATCGACACCAGCAAAGTGAATTATGGTGTAACGGTACTGCCGACCTTCAAGG
GTCAACCATCCAAACCGTTCGTTGGCGTGCTGAGCGCAGGTATTAACGCCGCCA
GTCCGAACAAAGAGCTGGCAAAAGAGTTCCTCGAAAACTATCTGCTGACTGATG
AAGGTCTGGAAGCGGTTAATAAAGACAAACCGCTGGGTGCCGTAGCGCTGAAGT
CTTACGAGGAAGAGTTGGCGAAAGATCCACGTATTGCCGCCACCATGGAAAACG
CCCAGAAAGGTGAAATCATGCCGAACATCCCGCAGATGTCCGCTTTCTGGTATGC
CGTGCGTACTGCGGTGATCAACGCCGCCAGCGGTCGTCAGACTGTCGATGAAGC
CCTGAAAGACGCGCAGACTAATTCGAGCTCCTCGAGC**TGA**

**6xHis-PhoCl1-MBP amino acid sequence**

MGSSHHHHHHSSGLVPRGSHMVIPDYFKQSFPEGYSWERSMTYEDGGICIATNDITM
EGDSFINKIHFKGTNFPPNGPVMQKRTVGWEASTEKMYERDGVLKGDVKMKLLLK
GGGHYRCDYRTTYKVKQKPVKLPDYHFVDHRIEILSHDKDYNKVKLYEHAVARNS
TDSMDELYKGGSGGMVSKGEETITSVIKPDMKNKLRMEGNVNGHAFVIEGEGSGKP
FEGIQTIDLEVKEGAPLPFAYDILTTAFHYGNRVFTKYPRKLGGGSGTMGIEEGKLVI
WINGDKGYNGLAEVGKKFEKDTGIKVTVEHPDKLEEKFPQVAATGDGPDIIFWAHD
RFGGYAQSGLLAEITPDKAFQDKLYPFTWDAVRYNGKLIAYPIAVEALSLIYNKDLL
PNPPKTWEEIPALDKELKAKGKSALMFNLQEPYFTWPLIAADGGYAFKYENGKYDI
KDVGVDNAGAKAGLTFLVDLIKNKHMNADTDYSIAEAAFNKGETAMTINGPWAWS
NIDTSKVNYGVTVLPTFKGQPSKPFVGVLSAGINAASPNKELAKEFLENYLLTDEGLE
AVNKDKPLGAVALKSYEEELAKDPRIAATMENAQKGEIMPNIPQMSAFWYAVRTA
VINAASGRQTVDEALKDAQTNSSSSS

**6xHis-PhoCl1-TEV nucleotide sequence**

**ATG**GGCAGCAGCCATCATCATCATCATCACAGCAGCGGCCTGGTGCCGCGCGGC
AGCCATATGGTGATCCCTGACTACTTCAAGCAGAGCTTCCCCGAGGGCTACAGCT
GGGAGCGCAGCATGACCTACGAGGACGGCGGCATCTGCATCGCCACCAACGACA
TCACAATGGAGGGGGACAGCTTCATCAACAAGATCCACTTCAAGGGCACGAACT
TCCCCCCCAACGGCCCCGTGATGCAGAAGAGGACCGTGGGCTGGGAGGCCAGCA
CCGAGAAGATGTACGAGCGCGACGGCGTGCTGAAGGGCGACGTGAAGATGAAG
CTGCTGCTGAAGGGCGGCGGCCACTATCGCTGCGACTACCGCACCACCTACAAG
GTCAAGCAGAAGCCCGTAAAGCTGCCCGACTACCACTTCGTGGACCACCGCATC
GAGATCCTGAGCCACGACAAGGACTACAACAAGGTGAAGCTGTACGAGCACGCC
GTGGCCCGCAACTCCACCGACAGCATGGACGAGCTGTACAAGGGTGGCAGCGGT
GGCATGGTGAGCAAGGGCGAGGAGACCATTACAAGCGTGATCAAGCCTGACATG
AAGAACAAGCTGCGCATGGAGGGCAACGTGAACGGCCACGCCTTCGTGATCGAG
GGCGAGGGCAGCGGCAAGCCCTTCGAGGGCATCCAGACGATTGATTTGGAGGTG

| |
|---|
| AAGGAGGGCGCCCCGCTGCCCTTCGCCTACGACATCCTGACCACCGCCTTCCACT<br>ACGGCAACCGCGTGTTCACCAAGTACCCACGGAAGCTTGGCGGCGGCTCTGGTA<br>CCGGATCCGGAGAAAGCTTGTTTAAGGGGCCGCGTGATTACAACCCGATATCGA<br>GCACCATTGTTCATTTGACGAATGAATCTGATGGGCACACAACATCGTTGTATGG<br>TATTGGATTTGGTCCCTTCATCATTACAAACAAGCACTTGTTTAGAAGAAATAAT<br>GGAACACTGGTGGTCCAATCACTACATGGTGTATTCAAGGTCAAGAACACCACG<br>ACTTTGCAACAACACCTCATTGATGGGAGGGACATGATAATTATTCGCATGCCTA<br>AGGATTTCCCACCATTTCCTCAAAAGCTGAAATTTAGAGAGCCACAAAGGGAAG<br>AGCGCATAGTCCTTGTGACAACCAACTTCCAAACTAAGAGCATGTCTAGCATGGT<br>GTCAGACACTAGTTCGACATTCCCTTCAGGAGATGGCATATTCTGGAAGCATTGG<br>ATTCAAACCAAGGATGGGCAGTGTGGCAGTCCATTAGTATCAACTAGAGATGGG<br>TTCATTGTTGGTATACACTCAGCATCGAATTTCACCAACACAAACAATTATTTCA<br>CAAGCGTGCCGAAAAACTTCATGGAATTGTTGACAAATCAGGAGGCGCAGCAGT<br>GGGTTAGTGGTTGGCGATTAAATGCTGACTCAGTATTGTGGGGAGGCCATAAAG<br>TTTTCATGGACAAACCTGAAGAGCCTTTTCAGCCAGTTAAGGAAGCGACTCAACT<br>CATGAATCGTCGTCGCCGTCGCTAA |
| **6xHis-PhoCl1-TEV amino acid sequence** |
| MGSSHHHHHHSSGLVPRGSHMVIPDYFKQSFPEGYSWERSMTYEDGGICIATNDITM<br>EGDSFINKIHFKGTNFPPNGPVMQKRTVGWEASTEKMYERDGVLKGDVKMKLLLK<br>GGGHYRCDYRTTYKVKQKPVKLPDYHFVDHRIEILSHDKDYNKVKLYEHAVARNS<br>TDSMDELYKGGSGGMVSKGEETITSVIKPDMKNKLRMEGNVNGHAFVIEGEGSGKP<br>FEGIQTIDLEVKEGAPLPFAYDILTTAFHYGNRVFTKYPRKLGGGSGTGSGESLFKGP<br>RDYNPISSTIVHLTNESDGHTTSLYGIGFGPFIITNKHLFRRNNGTLVVQSLHGVFKVK<br>NTTTLQQHLIDGRDMIIIRMPKDFPPFPQKLKFREPQREERIVLVTTNFQTKSMSSMVS<br>DTSSTFPSGDGIFWKHWIQTKDGQCGSPLVSTRDGFIVGIHSASNFTNTNNYFTSVPK<br>NFMELLTNQEAQQWVSGWRLNADSVLWGGHKVFMDKPEEPFQPVKEATQLMNRR<br>RRR |
| **6xHis-PhoCl1-V48ELP nucleotide sequence** |
| <u>**ATG**</u>GGCAGCAGCCATCATCATCATCATCACAGCAGCGGCCTGGTGCCGCGCGGC<br>AGCCATATGGTGATCCCTGACTACTTCAAGCAGAGCTTCCCCGAGGGCTACAGCT<br>GGGAGCGCAGCATGACCTACGAGGACGGCGGCATCTGCATCGCCACCAACGACA<br>TCACAATGGAGGGGGACAGCTTCATCAACAAGATCCACTTCAAGGGCACGAACT<br>TCCCCCCCAACGGCCCCGTGATGCAGAAGAGGACCGTGGGCTGGGAGGCCAGCA<br>CCGAGAAGATGTACGAGCGCGACGGCGTGCTGAAGGGCGACGTGAAGATGAAG<br>CTGCTGCTGAAGGGCGGCGGCCACTATCGCTGCGACTACCGCACCACCTACAAG<br>GTCAAGCAGAAGCCCGTAAAGCTGCCCGACTACCACTTCGTGGACCACCGCATC<br>GAGATCCTGAGCCACGACAAGGACTACAACAAGGTGAAGCTGTACGAGCACGCC<br>GTGGCCCGCAACTCCACCGACAGCATGGACGAGCTGTACAAGGGTGGCAGCGGT<br>GGCATGGTGAGCAAGGGCGAGGAGACCATTACAAGCGTGATCAAGCCTGACATG<br>AAGAACAAGCTGCGCATGGAGGGCAACGTGAACGGCCACGCCTTCGTGATCGAG<br>GGCGAGGGCAGCGGCAAGCCCTTCGAGGGCATCCAGACGATTGATTTGGAGGTG<br>AAGGAGGGCGCCCCGCTGCCCTTCGCCTACGACATCCTGACCACCGCCTTCCACT<br>ACGGCAACCGCGTGTTCACCAAGTACCCACGGAAGCTTGGCGGCGGCTCTGGGT<br>ACCGCAGCCATATGGGTGTTCCGGGCGTGGGTGTACCAGGTGTCGGTGTACCGG<br>GTGTCGGCGTACCTGGCGTCGGTGTCCCGGGTGTTGGTGTTCCGGGTGTAGGTGT<br>TCCGGGCGTGGGTGTACCAGGTGTCGGTGTACCGGGTGTCGGCGTACCTGGCGTC<br>GGTGTCCCGGGTGTTGGTGTTCCGGGTGTAGGTGTTCCGGGCGTGGGTGTACCAG<br>GTGTCGGTGTACCGGGTGTCGGCGTACCTGGCGTCGGTGTCCCGGGTGTTGGTGT<br>TCCGGGTGTAGGTGTTCCGGGCGTGGGTGTACCAGGTGTCGGTGTACCGGGTGTC<br>GGCGTACCTGGCGTCGGTGTCCCGGGTGTTGGTGTTCCGGGTGTAGGTGTTCCGG |

| |
|---|
| GCGTGGGTGTACCAGGTGTCGGTGTACCGGGTGTCGGCGTACCTGGCGTCGGTGT<br>CCCGGGTGTTGGTGTTCCGGGTGTAGGTGTTCCGGGCGTGGGTGTACCAGGTGTC<br>GGTGTACCGGGTGTCGGCGTACCTGGCGTCGGTGTCCCGGGTGTTGGTGTTCCGG<br>GTGTAGGTGTTCCGGGCGTGGGTGTACCAGGTGTCGGTGTACCGGGTGTCGGCGT<br>ACCTGGCGTCGGTGTCCCGGGTGTTGGTGTTCCGGGTGTAGGTGTTCCGGGCGTG<br>GGTGTACCAGGTGTCGGTGTACCGGGTGTCGGCGTACCTGGCGTCGGTGTCCCGG<br>GTGTTGGTGTTCCGGGTGTAGGTTAC**TGA** |
| **6xHis-PhoCl1-V48ELP amino acid sequence** |
| MGSSHHHHHHSSGLVPRGSHMVIPDYFKQSFPEGYSWERSMTYEDGGICIATNDITM<br>EGDSFINKIHFKGTNFPPNGPVMQKRTVGWEASTEKMYERDGVLKGDVKMKLLLK<br>GGGHYRCDYRTTYKVKQKPVKLPDYHFVDHRIEILSHDKDYNKVKLYEHAVARNS<br>TDSMDELYKGGSGGMVSKGEETITSVIKPDMKNKLRMEGNVNGHAFVIEGEGSGKP<br>FEGIQTIDLEVKEGAPLPFAYDILTTAFHYGNRVFTKYPRKLGGGSGYRSHMGVPGV<br>GVPGVGVPGVGVPGVGVPGVGVPGVGVPGVGVPGVGVPGVGVPGVGVPGVGVPG<br>VGVPGVGVPGVGVPGVGVPGVGVPGVGVPGVGVPGVGVPGVGVPGVGVPGVGVP<br>GVGVPGVGVPGVGVPGVGVPGVGVPGVGVPGVGVPGVGVPGVGVPGVGVPGVGV<br>PGVGVPGVGVPGVGVPGVGVPGVGVPGVGVPGVGVPGVGVPGVGVPGVGVPGVG<br>VPGVGVPGVGVPGVGVPGVGY |
| **CTPF-R5 amino acid sequence** |
| HYGNRVFTKYPRKLGGGSSSSKKSGSYSGSKGSKRRI |
| **CTPF-*Mj*RibK amino acid sequence** |
| HYGNRVFTKYPRKLGGGSLVKLMIIEGEVVSGLGEGRYFLSLPPYKEIFKKILGF<br>EPYEGTLNLKLDREFDINKFKYIETEDFEFNGKRFFGVKVLPIKILIGNKKIDGAIVVP<br>KKTYHSSEIIEIIAPMKLREQFNLKDGDVIKILIKGDKDE |
| **CTPF-LBT amino acid sequence** |
| HYGNRVFTKYPRGGGGSDPDKDGTIDLKE |
| **CTPF-MBP amino acid sequence** |
| HYGNRVFTKYPRKLGGGSGTMGIEEGKLVIWINGDKGYNGLAEVGKKFEKDTGIKV<br>TVEHPDKLEEKFPQVAATGDGPDIIFWAHDRFGGYAQSGLLAEITPDKAFQDKLYPF<br>TWDAVRYNGKLIAYPIAVEALSLIYNKDLLPNPPKTWEEIPALDKELKAKGKSALMF<br>NLQEPYFTWPLIAADGGYAFKYENGKYDIKDVGVDNAGAKAGLTFLVDLIKNKHM<br>NADTDYSIAEAAFNKGETAMTINGPWAWSNIDTSKVNYGVTVLPTFKGQPSKPFVG<br>VLSAGINAASPNKELAKEFLENYLLTDEGLEAVNKDKPLGAVALKSYEEELAKDPRI<br>AATMENAQKGEIMPNIPQMSAFWYAVRTAVINAASGRQTVDEALKDAQTNSSSSS |
| **CTPF-TEV amino acid sequence** |
| HYGNRVFTKYPRKLGGGSGTGSGESLFKGPRDYNPISSTIVHLTNESDGHTTSLYGIG<br>FGPFIITNKHLFRRNNGTLVVQSLHGVFKVKNTTTLQQHLIDGRDMIIIRMPKDFPPFP<br>QKLKFREPQREERIVLVTTNFQTKSMSSMVSDTSSTFPSGDGIFWKHWIQTKDGQCG<br>SPLVSTRDGFIVGIHSASNFTNTNNYFTSVPKNFMELLTNQEAQQWVSGWRLNADSV<br>LWGGHKVFMDKPEEPFQPVKEATQLMNRRRRR |
| **CTPF-V48ELP or CTPF-ELP amino acid sequence** |
| HYGNRVFTKYPRKLGGGSGYRSHMGVPGVGVPGVGVPGVGVPGVGVPGVGVPGV<br>GVPGVGVPGVGVPGVGVPGVGVPGVGVPGVGVPGVGVPGVGVPGVGVPGVGVPG<br>VGVPGVGVPGVGVPGVGVPGVGVPGVGVPGVGVPGVGVPGVGVPGVGVPGVGVP<br>GVGVPGVGVPGVGVPGVGVPGVGVPGVGVPGVGVPGVGVPGVGVPGVGVPGVGV<br>PGVGVPGVGVPGVGVPGVGVPGVGVPGVGVPGVGVPGVGVPGVGVPGVGY |

**Table S2.** Primers used to make the gene products of PhoCl1-R5, PhoCl1-LBT, *Mj*RibK, and supercharged proteins. The forward and reverse primers have been indicated by the suffixes fwd and rev respectively.

| # | Primers | 5'-3' sequence |
|---|---|---|
| **For construction of 6xHis-PhoCl1-R5 and 6xHis-PhoCl1-LBT from 6xHis-PhoCl1 (by RF Cloning)** | | |
| 1 | PhoCl1-R5-RF-fwd1 | ACAAGCGTGATCAAGCCTGAC |
| 2 | PhoCl1-R5-RF-rev1 | GTGCGGCCGCAAGCTTTTACAGGATGCGACGTTTGGAGCCTTTGCTACCAGAGTAGGAACCAGACTTTTAGAGGAAGAAGAGCCGCCGCCAAGCTTCCGTGGGTACTTGGTGAACAC |
| 3 | PhoCl1-LBT-RF-fwd1 | ACAAGCGTGATCAAGCCTGAC |
| 4 | PhoCl1-LBT-RF-rev1 | GTGCGGCCGCAAGCTTTTATTCTTTCAGATCAATGGTGCCATCTTTATCCGGATCGGAGCCGCCGCCGCCCCGTGGGTACTTGGTGAACAC |
| **For construction of 6xHis-PhoCl1-Linker from 6xHis-PhoCl1 (by RF Cloning)** | | |
| 1 | PhoCl1-Linker-fwd1 | ACAAGCGTGATCAAGCCTGAC |
| 2 | PhoCl1-Linker-rev1 | CTTTGTTAGCAGCCGGATCTCTCAGCTCGAGGAGCTCGGATCCGGTACCAGAGCCGCCGCCAAGCTTCCGTGGGTACTTGGTGAACA |
| **For fusion construct PhoCl1-MjRibK (by Hi-Fi Cloning)** | | |
| 1 | PhoCl1_fwd | GTTTAACTTTAAGAAGGAGATATACCATGGGCAGCAGCCATCATCATCATCATCACGGCGGTACCGTGATCCCTGACTACTTCAAGC |
| 2 | PhoCl1_rev | AGAGCCGCCGCCAAGCTTCCGTGGGTACTTGGTGAAC |
| 3 | *Mj*RibK_fwd | CGGAAGCTTGGCGGCGGCTCTTTGGTGAAATTGATGATTATTG |
| 4 | *Mj*RibK_rev | TCAGTGGTGGTGGTGGTGGTGCTCGAGTTATTCATCTTTATCTCCCTTAAT |
| **For fusion construct PhoCl1-MBP and PhoCl1-TEV the MBP-TEV and PhoCl1 is added with required restriction sites (by Hi-Fi Cloning)** | | |
| 1 | MBP_fwd | GAATTGTGAGCGGATAACAATCATCATCATCATCATCATGGTACCATGGGAATCGAAGAAGGT |
| 2 | MBP_rev | CCAGATTGAGCGTAGCCAC |
| 3 | TEV_fwd | AGGGGAGATTTTGCAGGTGGATCCGGAGAAAGCTTGTTTAAGGGG |

| | | |
|---|---|---|
| 4 | TEV_rev | GGGAAATCCTTAGGCATGCGAATAAT |
| 5 | PhoCl1-MBPTEV-Fwd | ACAAGCGTGATCAAGCCTGAC |
| 6 | PhoCl1-MBPTEV-Rev | CTTTGTTAGCAGCCGGATCTCTCAGCTCGAGGAGCTCGGATCCGGTACCAGAGCCGCCGCCAAGCTTCCGTGGGTACTTGGTGAACA |

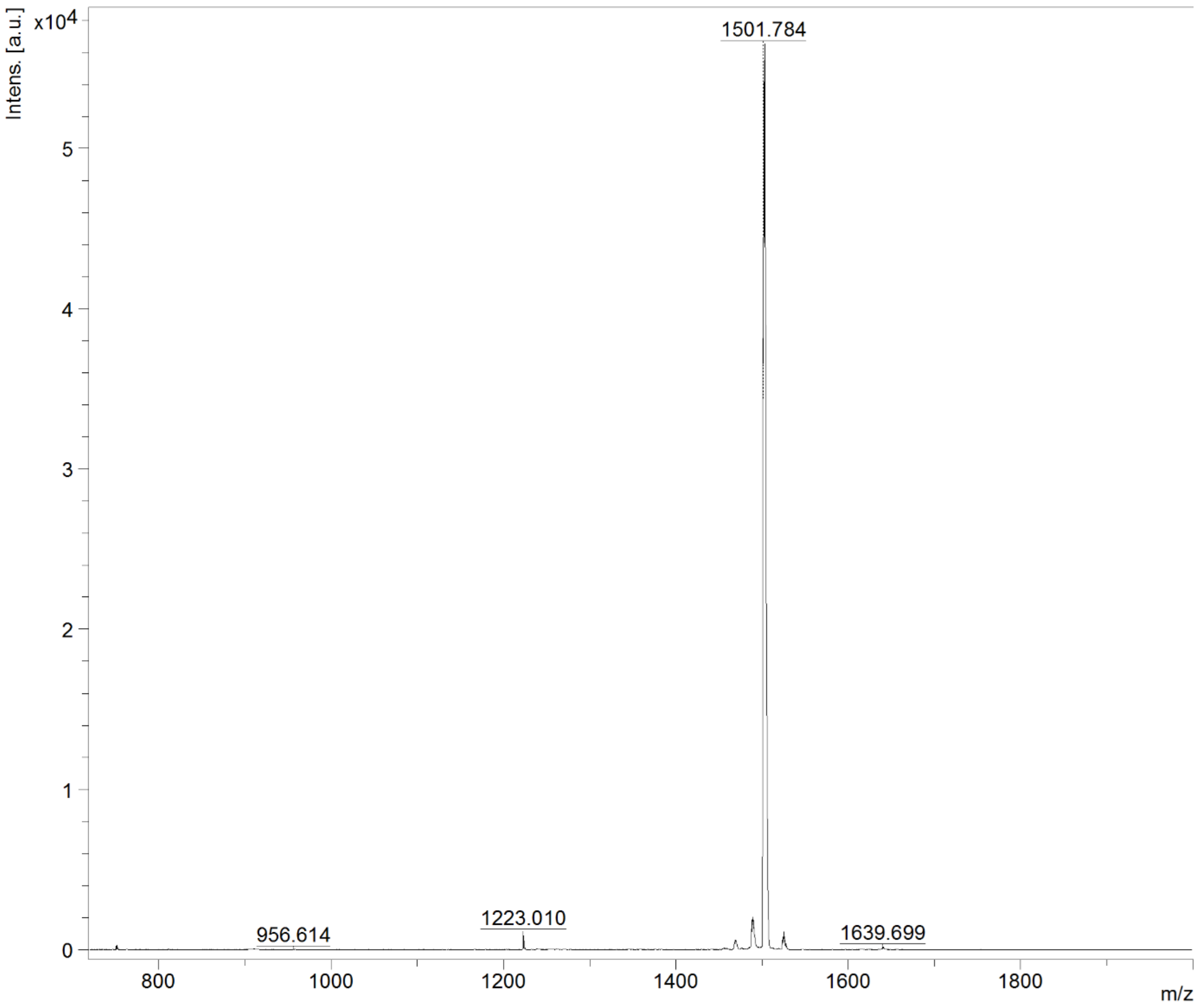


**Figure S1. MALDI-TOF mass spectrum of purified CTPF in the range of m/z 800 to 2000.** There is a prominent m/z peak at 1501.784, which is close to the theoretically predicted MW of CTPF (1500.688 Da).

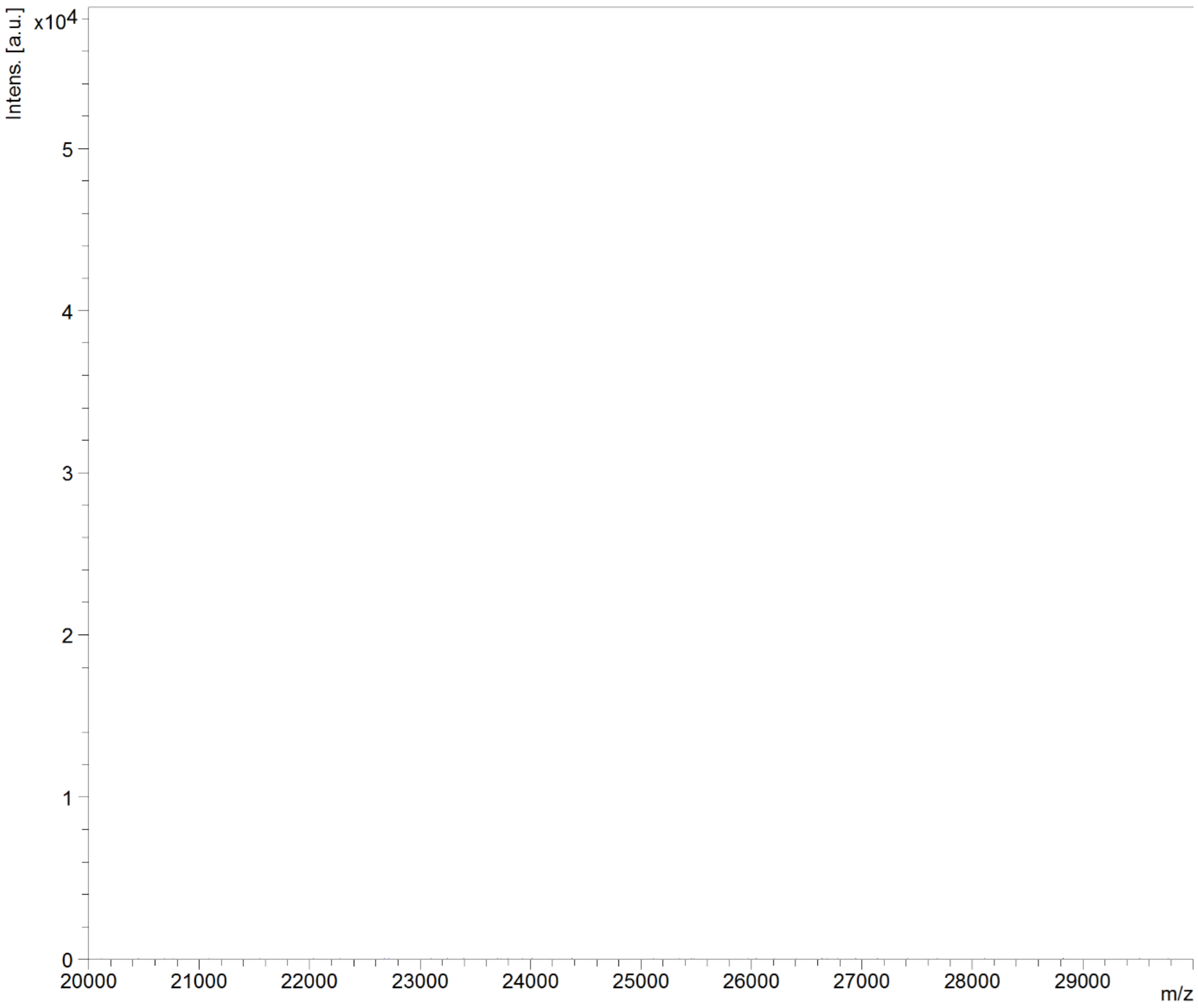


**Figure S2. MALDI-TOF mass spectrum of purified CTPF in the range of m/z 20000-30000.** Absence of any significant peak in this region indicates the purity of CTPF and rules out the possibility of PhoCl1 and PhoCl1 empty barrel contamination.

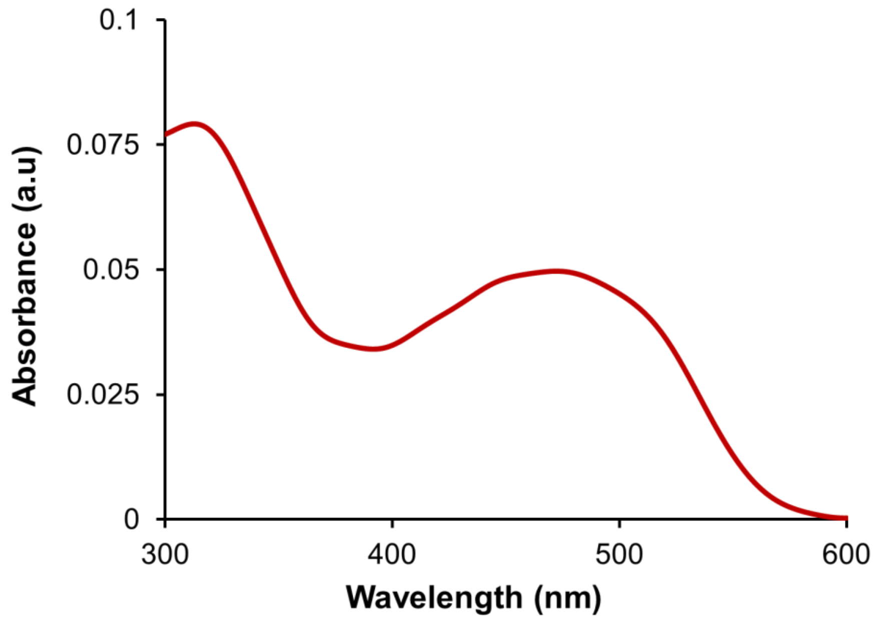


**Figure S3.** Absorbance Spectra of CTPF, showing 2 peaks, one at 320 nm and the other at 470 nm.

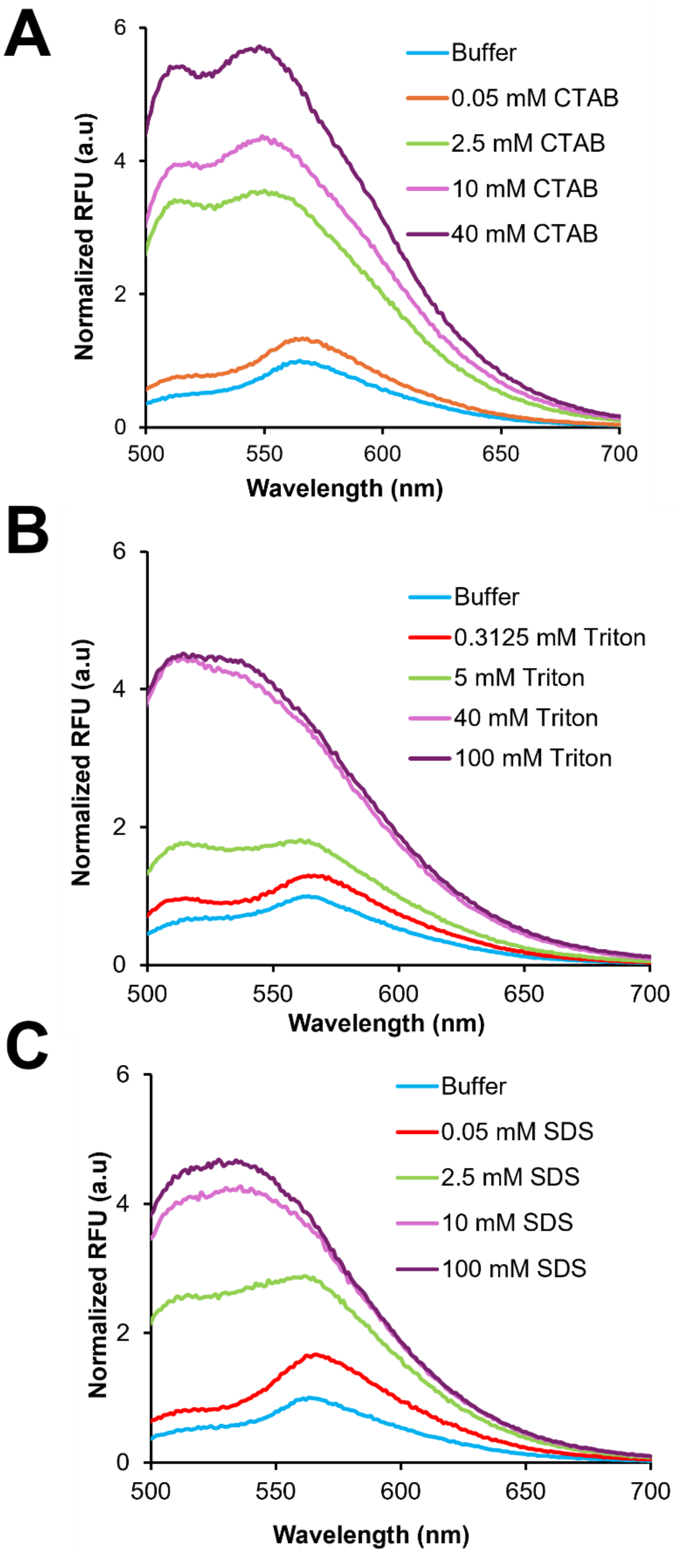


**Figure S4**. **Spectral studies of CTPF in various detergents. (A)** Normalized fluorescence emission spectra of CTPF at 470 nm when subjected to different concentrations of CTAB (Cetyltrimethylammonium Bromide), **(B)** Normalized fluorescence emission spectra of CTPF at 470 nm when subjected to different concentrations of Triton X-100, and (**C**) Normalized fluorescence emission spectra of CTPF at 470 nm when subjected to different concentrations of SDS (Sodium Dodecyl Sulfate).

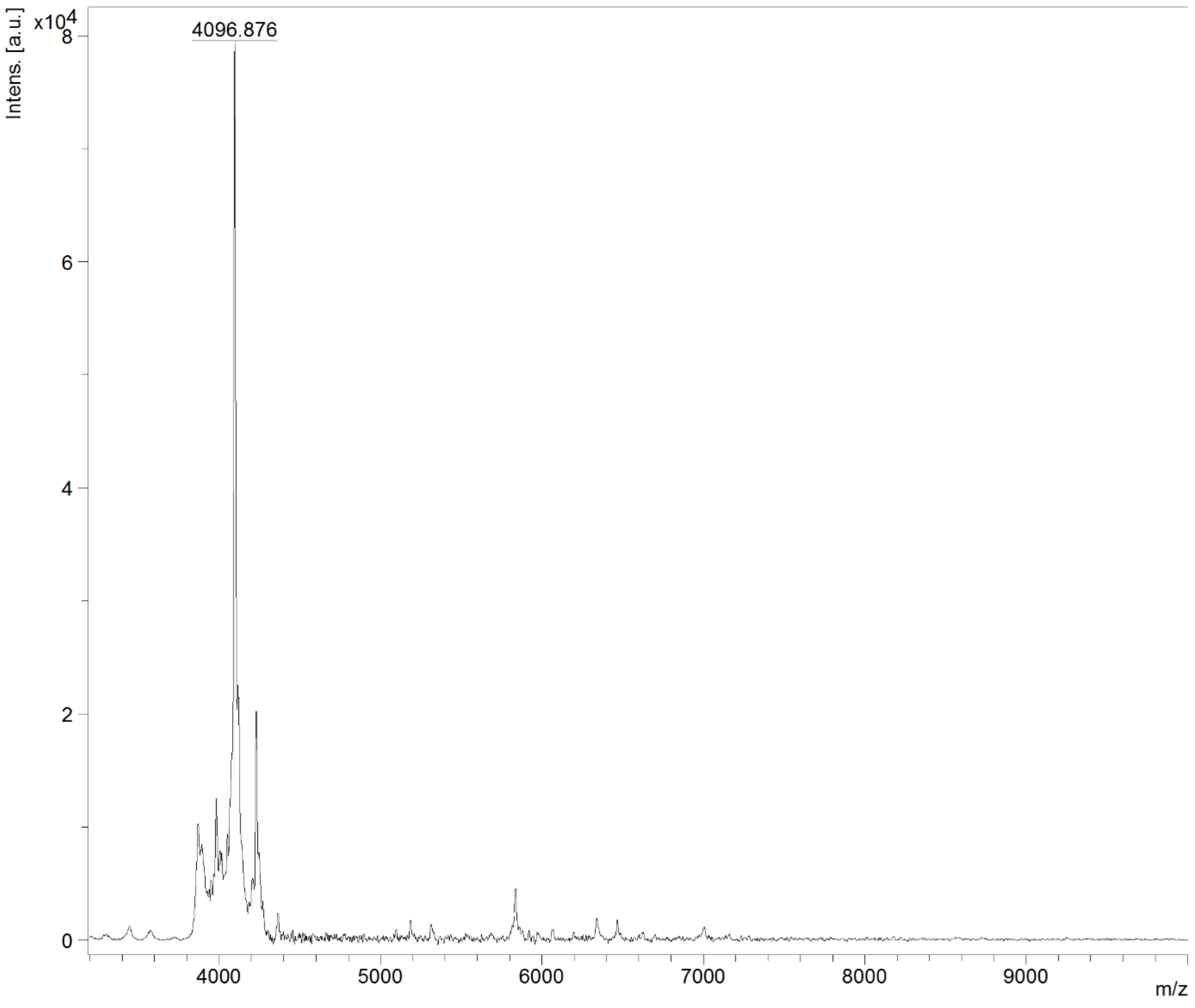


**Figure S5. MALDI-TOF mass spectrum of purified CTPF-R5 in the range of m/z 3500 to 10000.** There is a prominent m/z peak at 4096.87, which is close to theoretically predicted MW of CTPF-R5 (4082.62 Da).

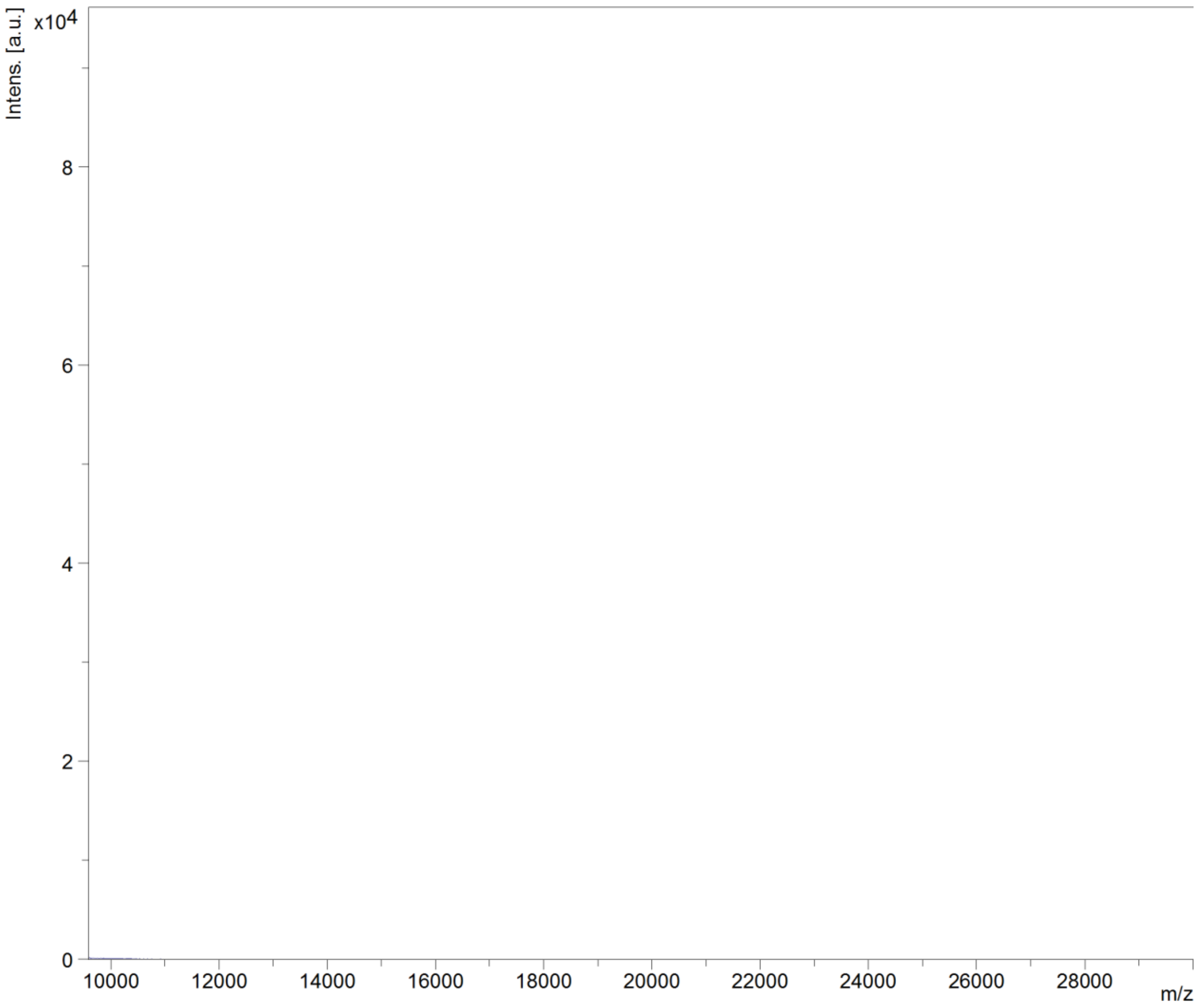


**Figure S6. MALDI-TOF mass spectrum of purified CTPF-R5 in the range of m/z 10000-30000.** Absence of any significant peak in this region indicates the purity of R5 and discards the possibility of PhoCl1 barrel contamination.

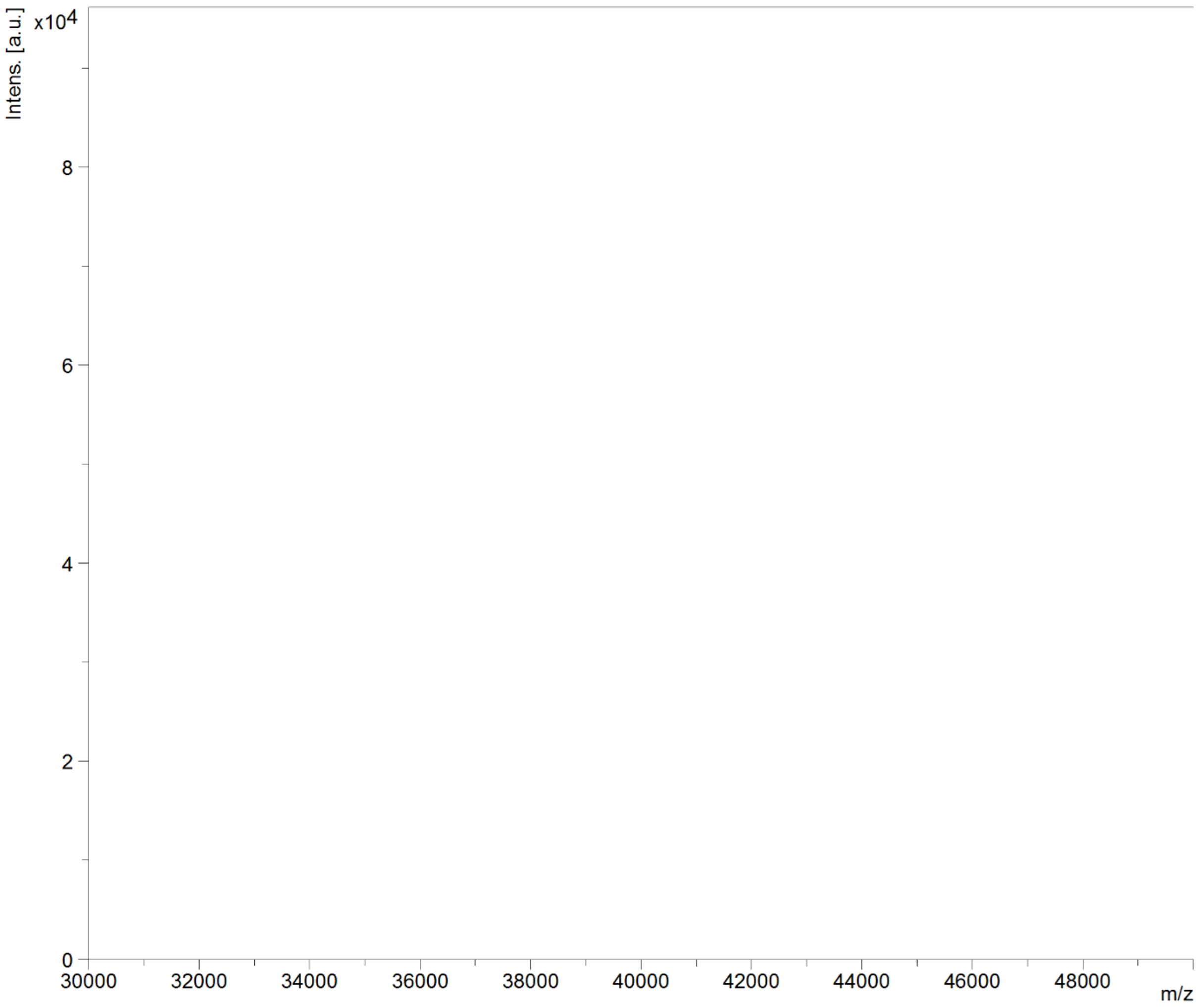


**Figure S7. MALDI-TOF Mass Spectrum of purified CTPF-R5 in the range of m/z 30000-50000.** Absence of any significant peak in this region indicates the purity of R5 and discards the possibility of fusion protein contamination.

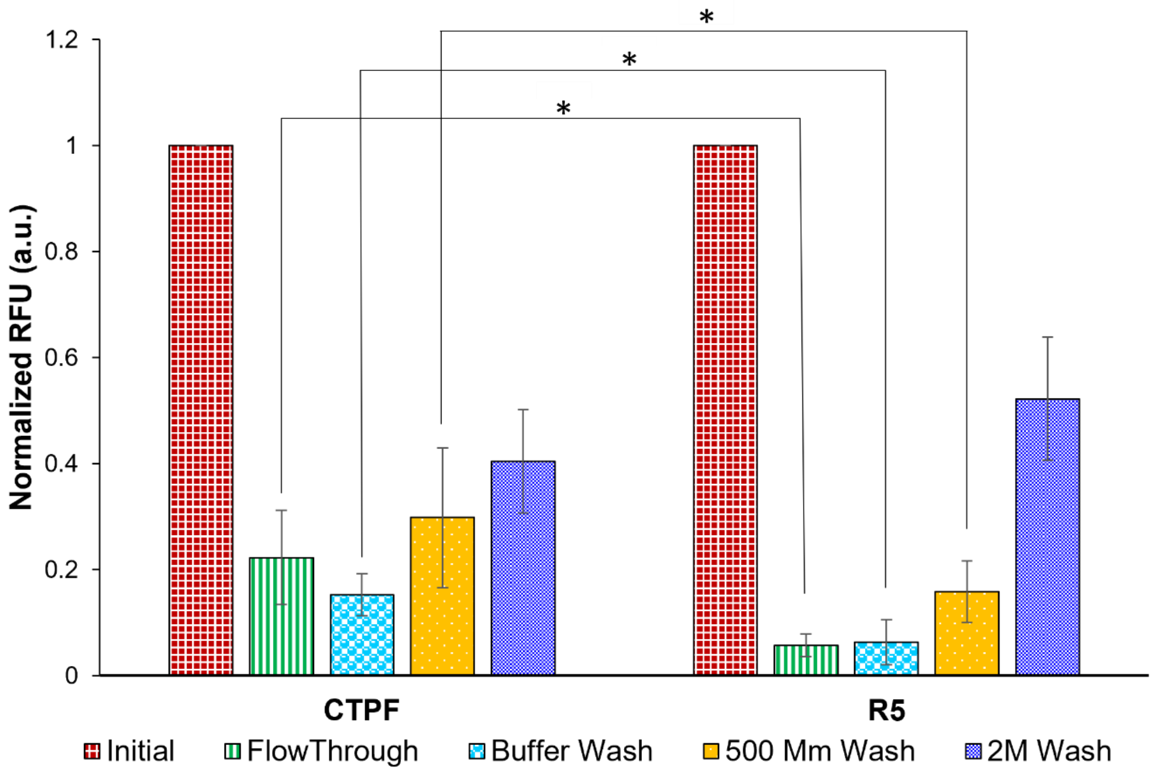


**Figure S8. Adsorption of CTPF-R5 on silica beads compared to CTPF.** Compared to the initial fluorescence (red filled squares), the fluorescence in the flow-through (green filled lines) of both CTPF-R5 and CTPF decreased over 30 minutes. During the buffer wash (light blue filled circles), significantly more loosely bound CTPF was released from the beads compared to CTPF-R5. Similarly, CTPF was eluted to a greater extent than CTPF-R5 during the 500 mM lysine wash (yellow background), indicating weaker binding affinity. Finally, the 2 M wash (blue filled squares) eluted all remaining bound material.

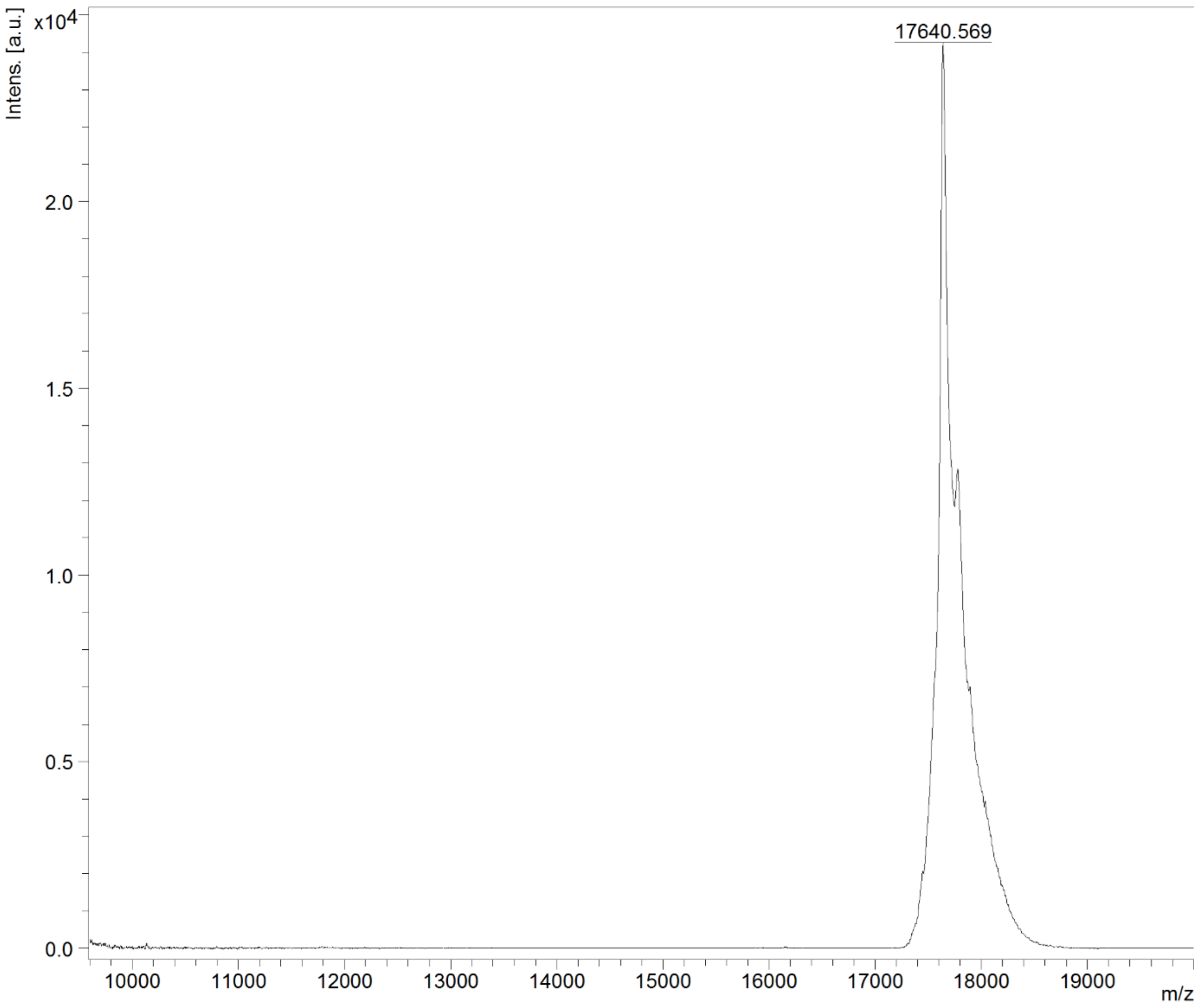


**Figure S9. MALDI-TOF Mass spectrum of purified CTPF-*Mj*RibK in the range of m/z 10000-20000.** There is a prominent m/z peak at 17640.569, which is close to the theoretically predicted MW of CTPF-*Mj*RibK 17649.688 Da.

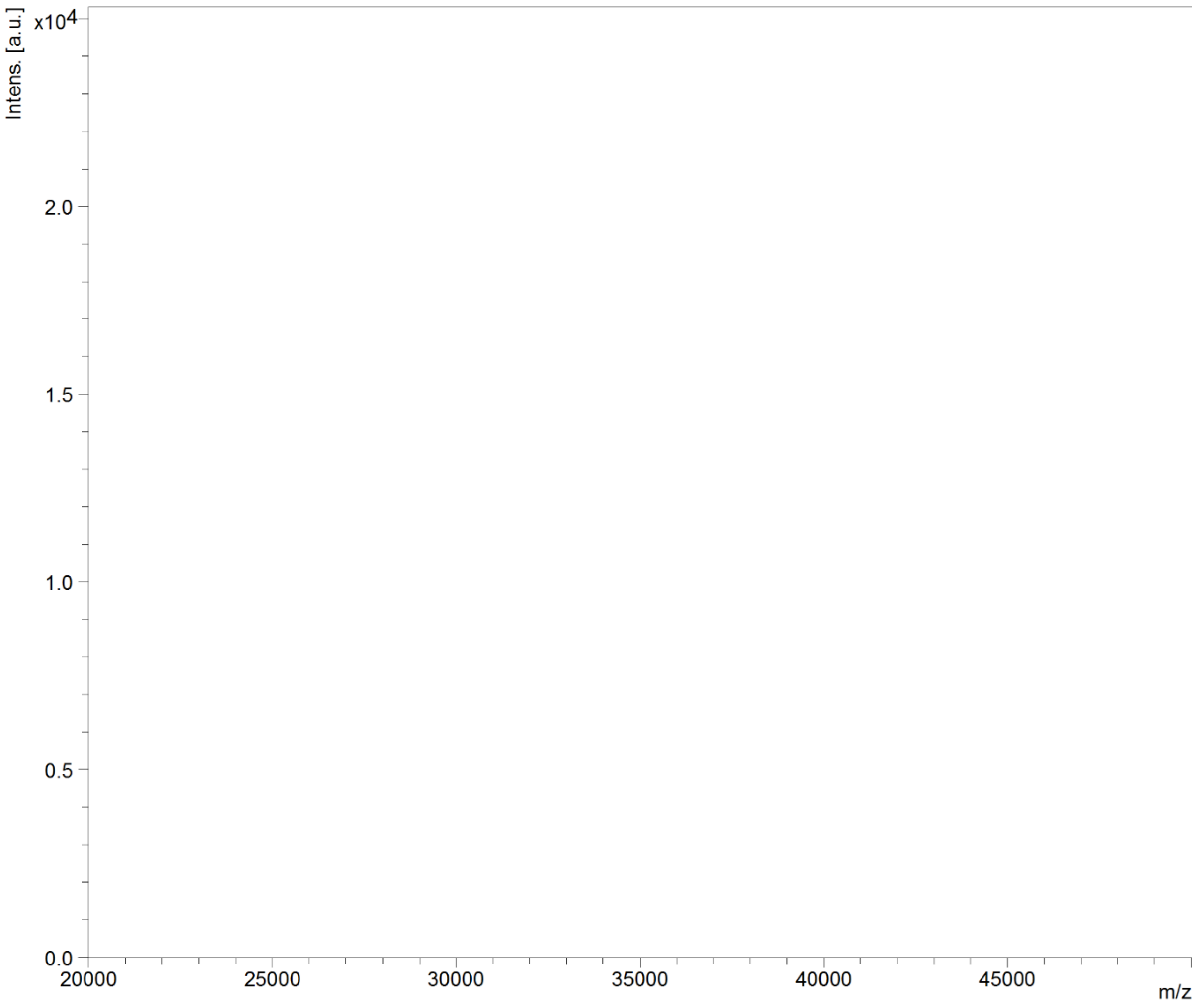


**Figure S10. MALDI-TOF Mass spectrum of purified CTPF-*Mj*RibK in the range of m/z 20000-50000.** Absence of any significant peak in this region indicates the purity of *Mj*RibK and discards the possibility of PhoCl1 empty barrel and fusion protein contamination.

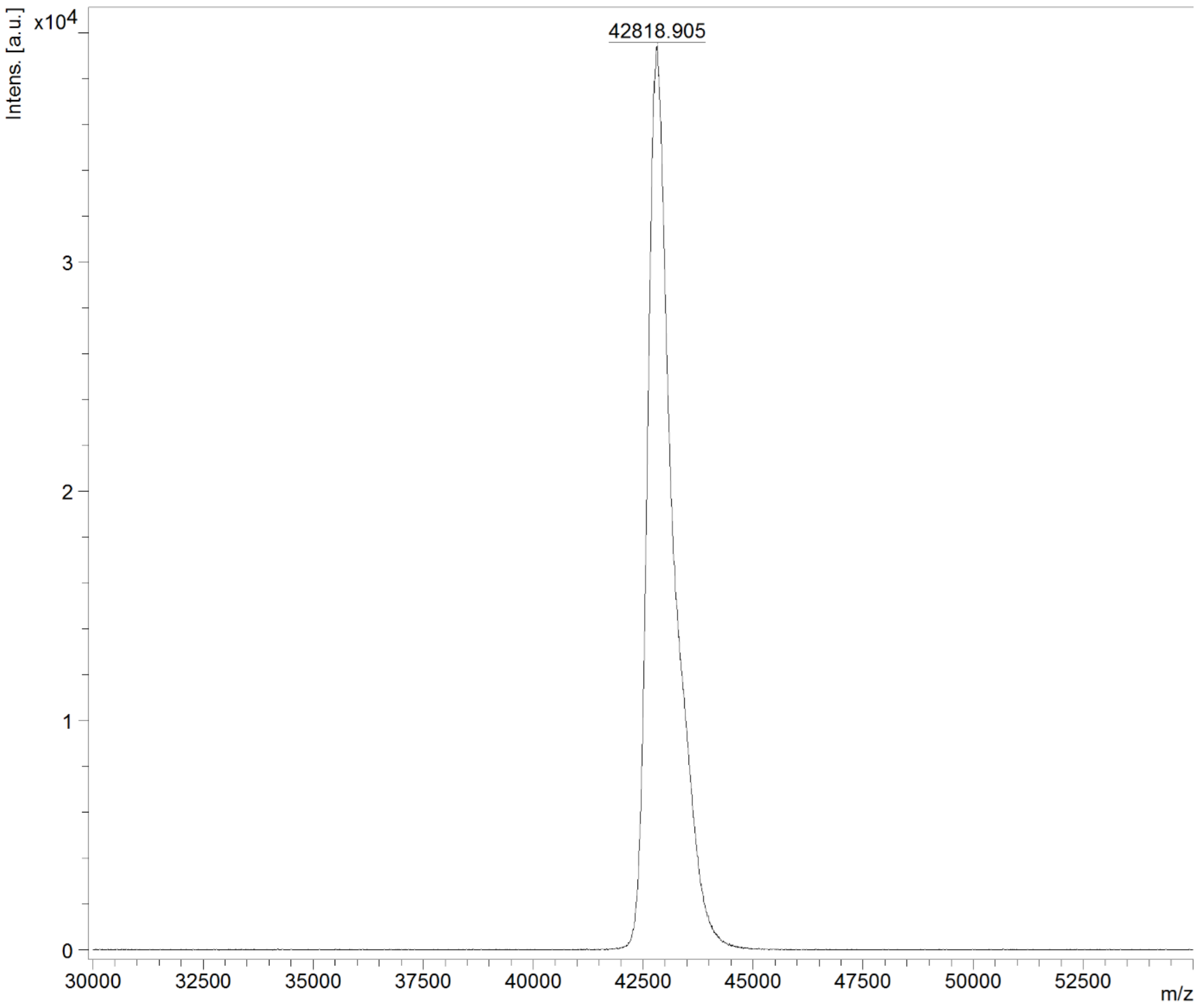


**Figure S11. MALDI-TOF Mass spectrum of purified CTPF-MBP in the range of m/z 30000 to 53000.** There is a prominent m/z peak at 42818.905, which is close to the theoretically predicted MW of CTPF-MBP 42958.67 Da

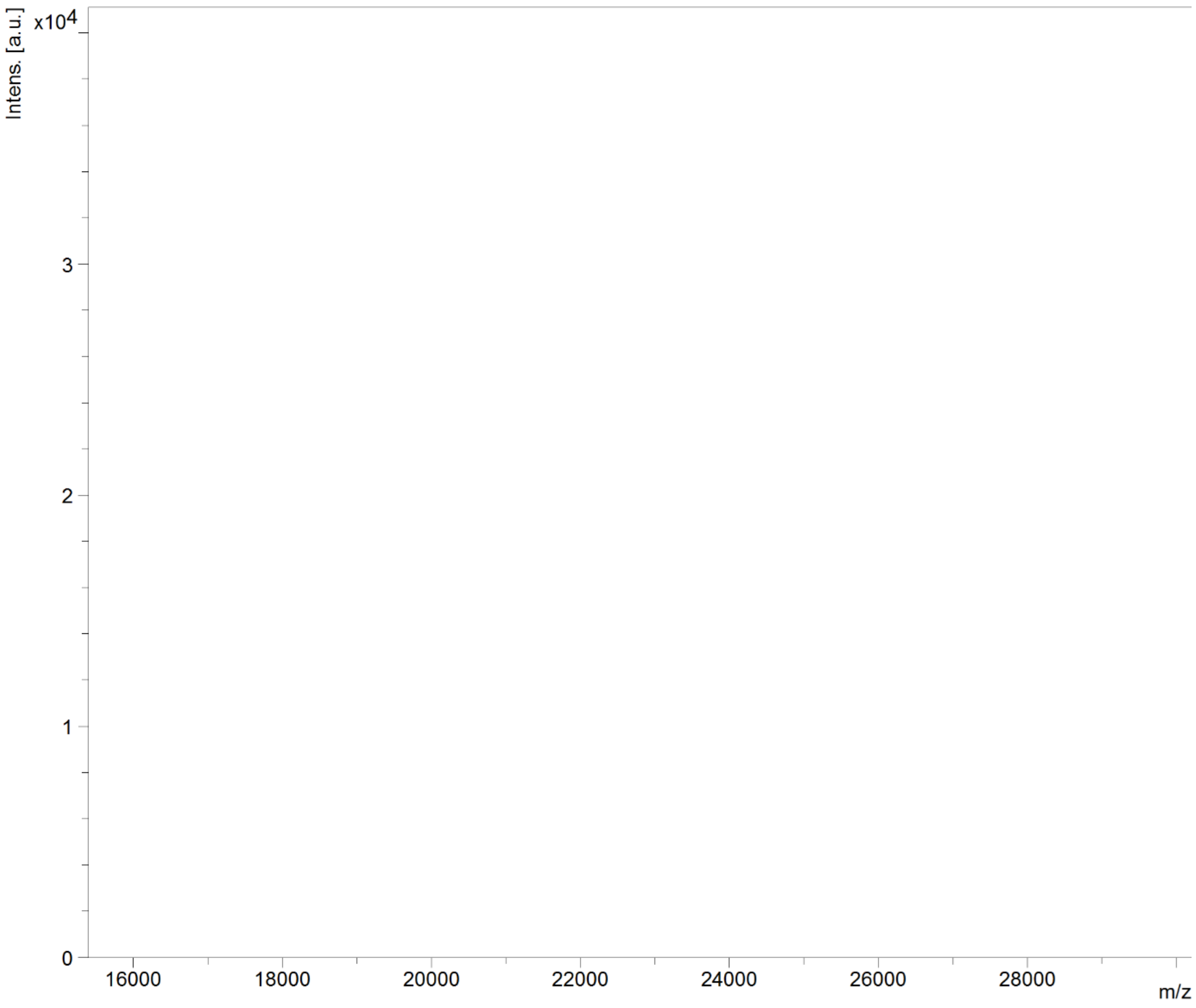


**Figure S12. MALDI-TOF Mass spectrum of purified CTPF-MBP in the range of m/z 16000-30000.** Absence of any significant peak in this region indicates the purity of MBP and discards the possibility of PhoCl1 barrel contamination.

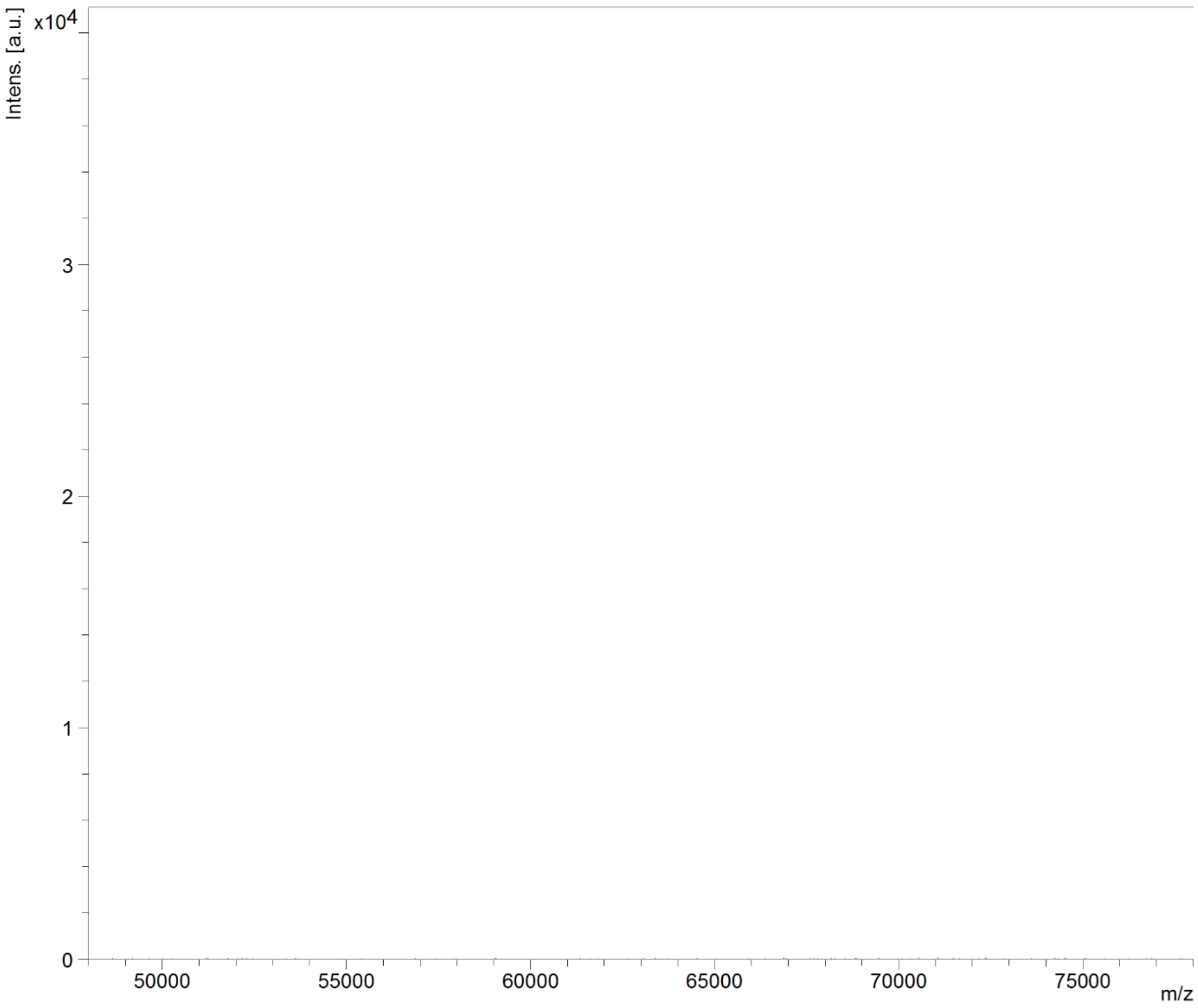


**Figure S13. MALDI-TOF Mass spectrum of purified CTPF-MBP in the range of m/z 50000-75000.** Absence of any significant peak in this region indicates the purity of MBP and discards the possibility of fusion protein contamination.

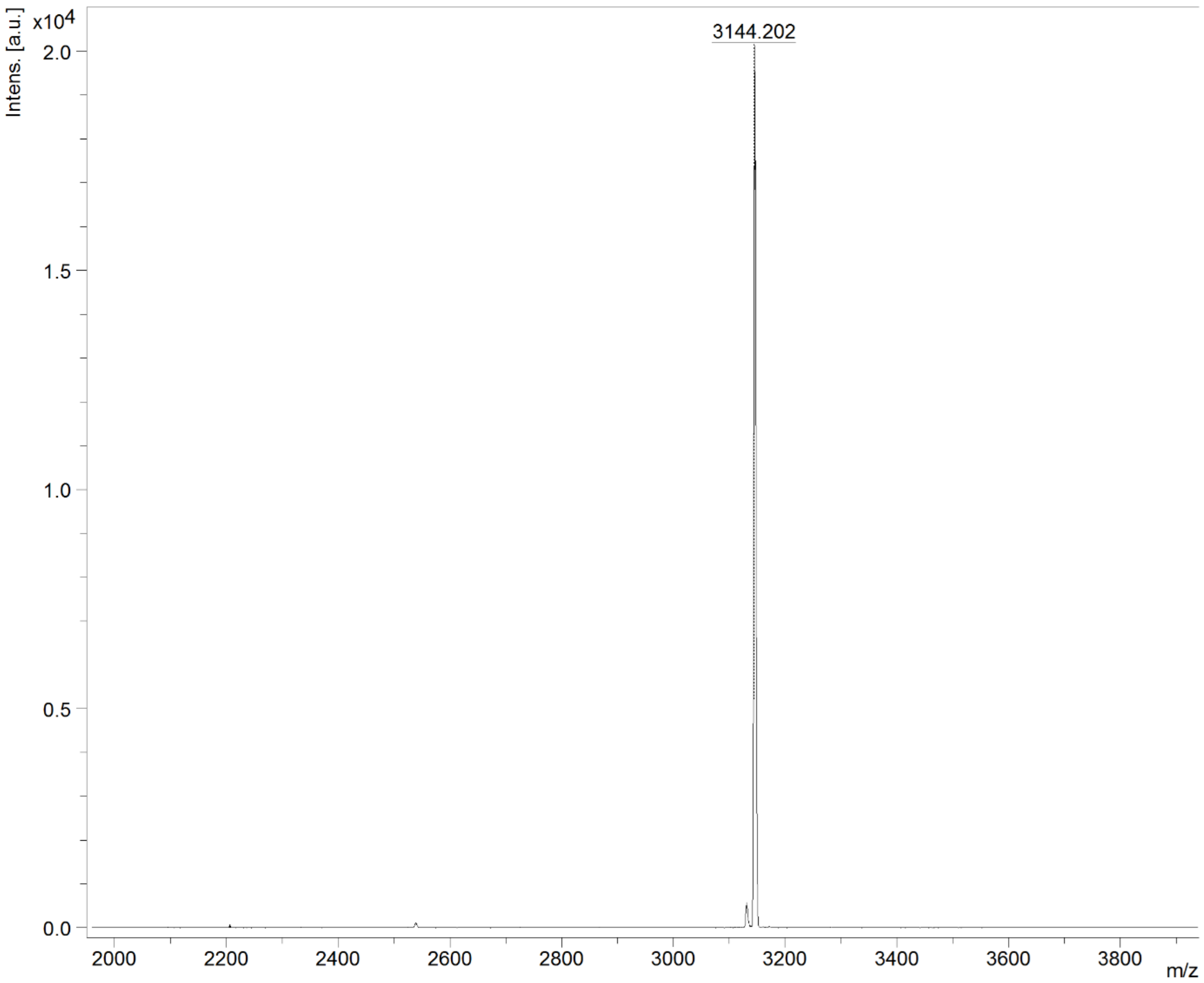


**Figure S14. MALDI-TOF Mass spectrum of purified CTPF-LBT in the range of m/z 2000 to 4000.** There is a prominent m/z peak at 3144.202, which is close to the theoretically predicted Mw of CTPF-LBT 3143.385 Da.

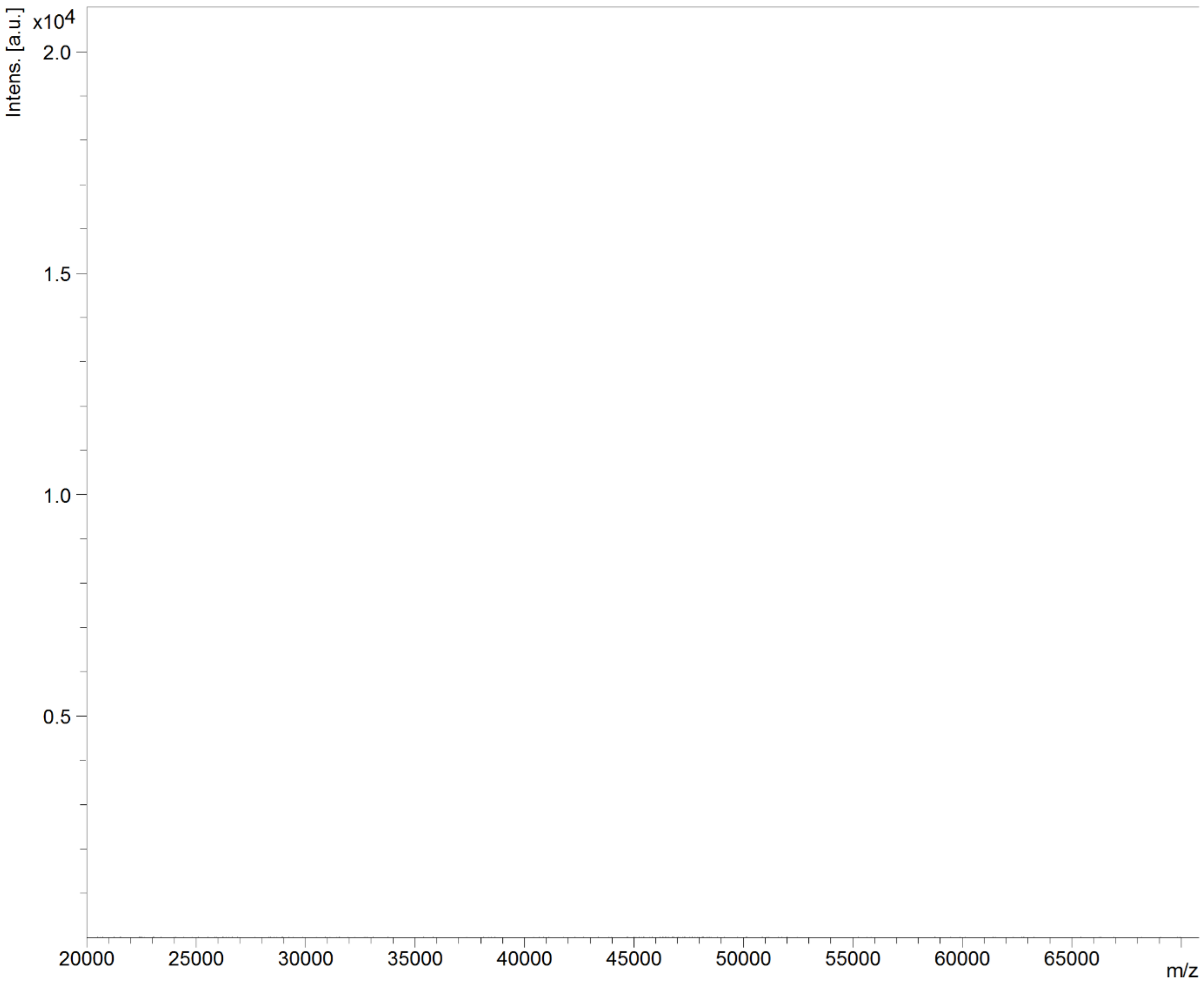


**Figure S15. MALDI-TOF Mass spectrum of purified CTPF-LBT in the range of m/z 20000-70000.** Absence of any significant peak in this region indicates the purity of CTPF-LBT and discards the possibility of PhoCl1-empty barrel protein contamination.

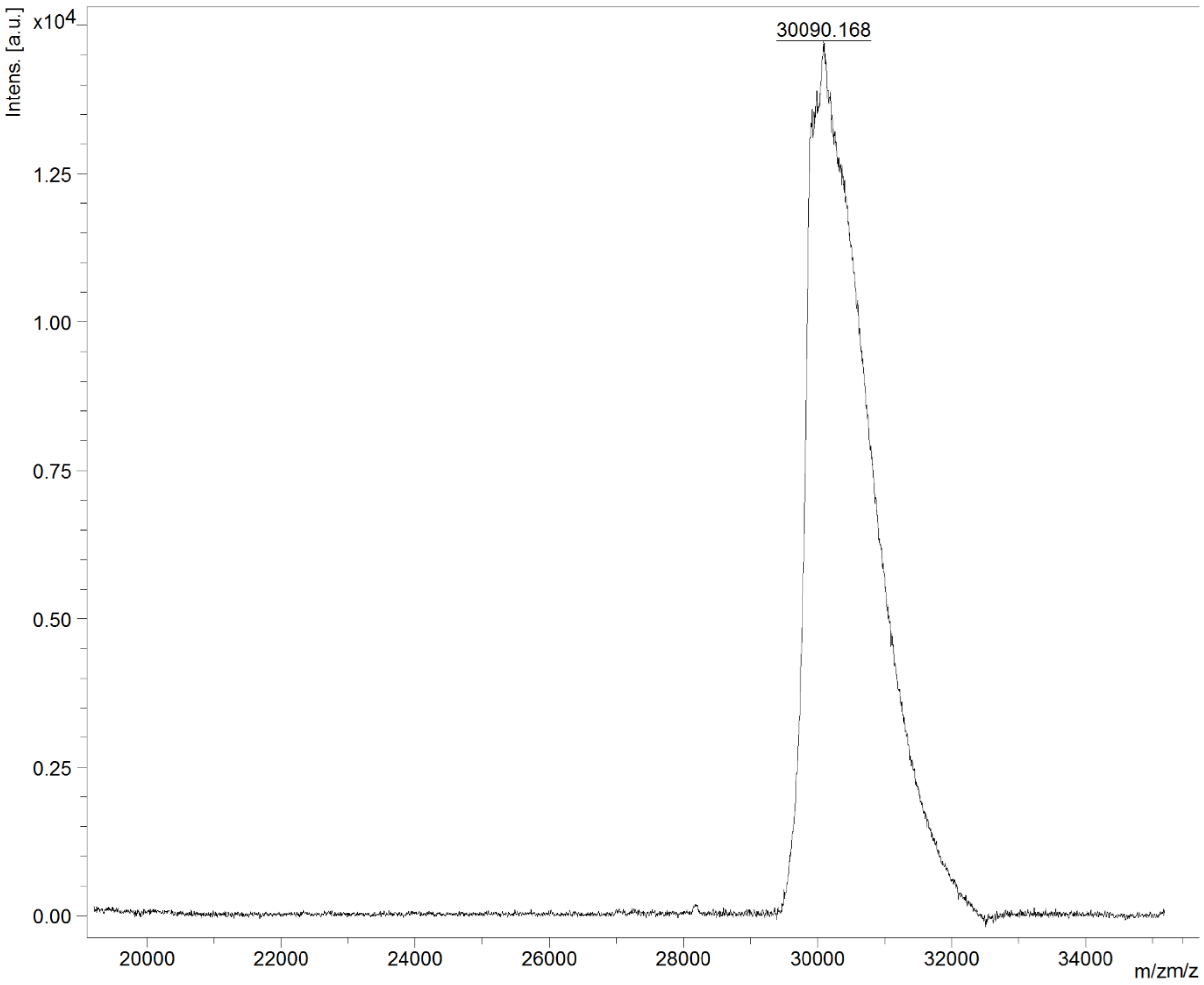


**Figure S16. MALDI-TOF Mass spectrum of purified CTPF-TEV in the range of m/z 20000 to 35000.** There is a prominent m/z peak at 30090.168, which is close to the theoretically predicted Mw of CTPF-TEV 29832.83 Da

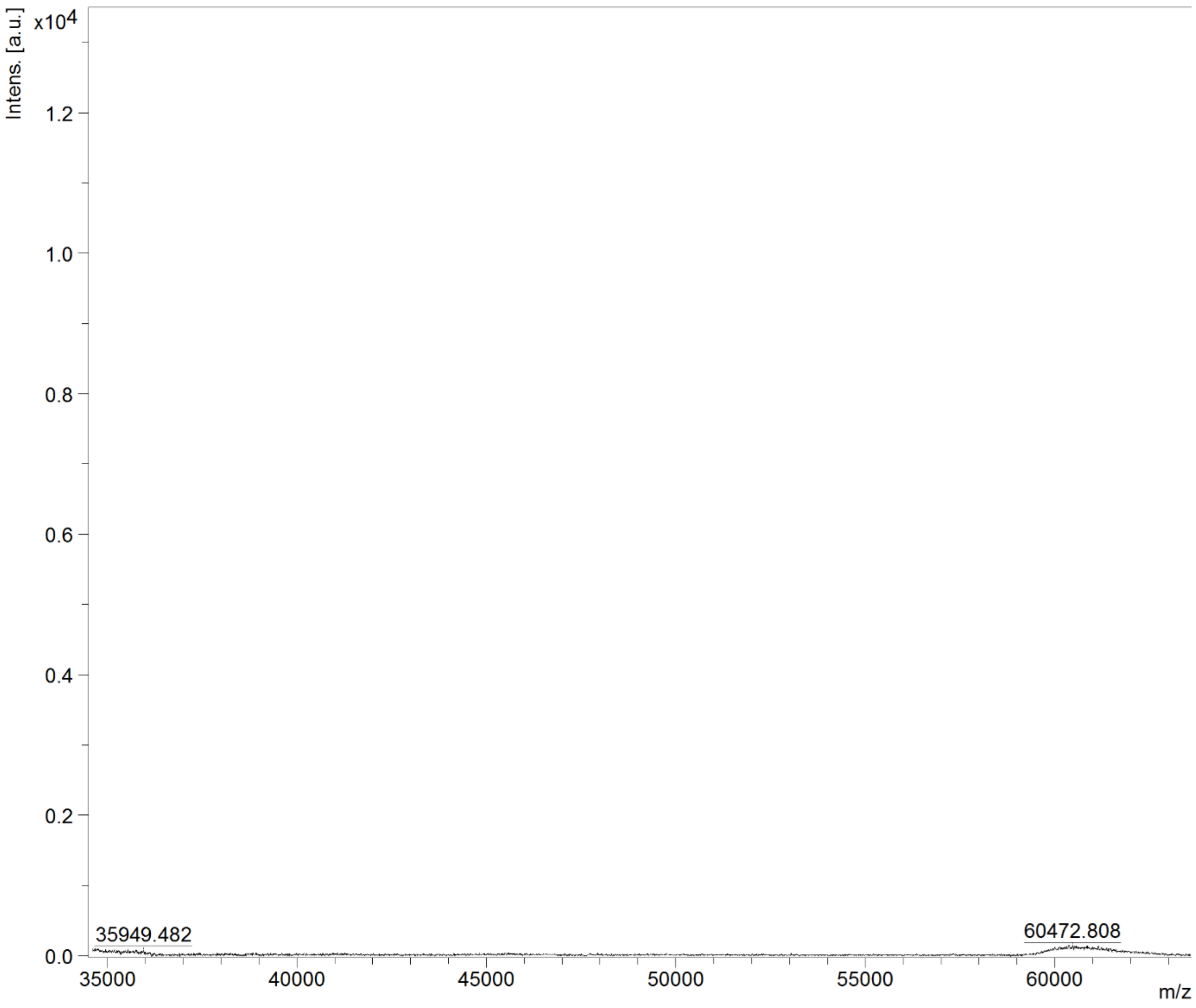


**Figure S17. MALDI-TOF Mass spectrum of purified CTPF-TEV in the range of m/z 35000-60000.** Absence of any significant peak in this region indicates the purity of CTPF-TEV and discards the possibility of fusion protein contamination.

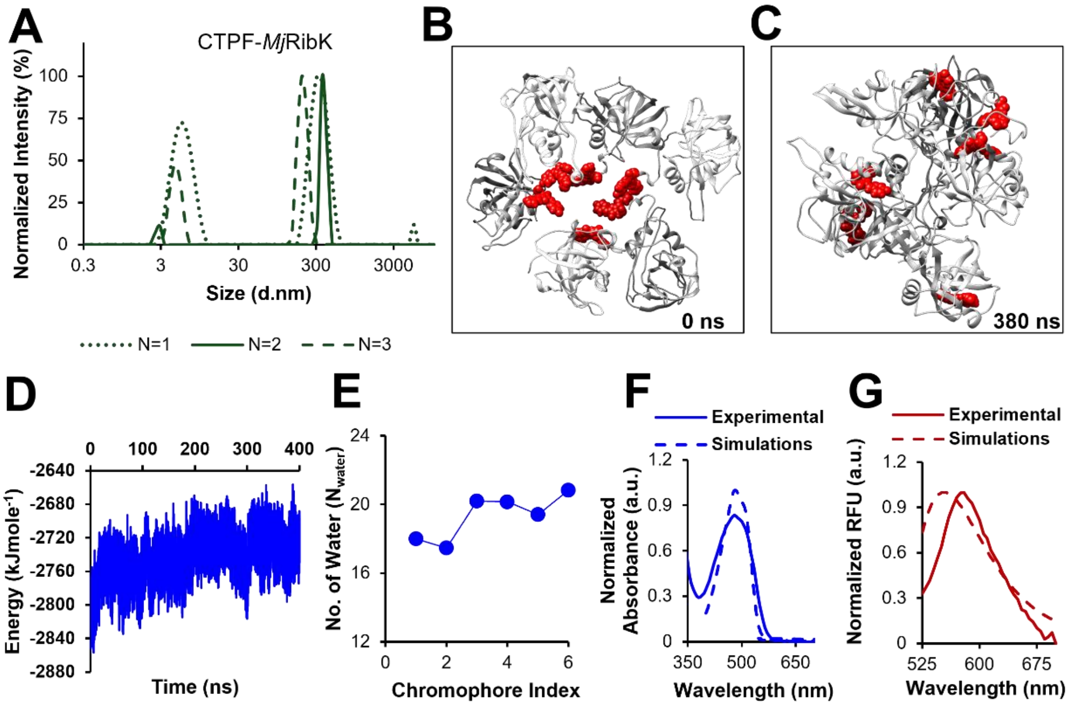


**Figure S18. Size Analysis of *in situ* generated CTPF-*Mj*RibK and computational validation of red fluorescence observation.** (A) CTPF-*Mj*RibK showed 2 prominent peaks, 4 ± 1.5 nm, and 289 ± 84 nm (major population), **(B)** Molecular dynamics snapshots of chromophore aggregation at time 0 ns, **(C)** Molecular dynamics snapshots of chromophore aggregation at time 380 ns. The chromophore is shown in red using a van der Waals representation, while the protein~~eptide~~ is represented in gray, **(D)** Change in total interaction energy depending on time among all chromophores in molecular dynamics trajectory, **(E)** Number of water molecules (hydration number) around chromophores in 6 CTPF-*Mj*RibK molecules, **(F)** Absorbance and **(G)** emission scan acquired through QM/MM (dashed lines) by considering hydration number – overlayed with experimentally observed scans (solid lines) of CTPF-*Mj*RibK

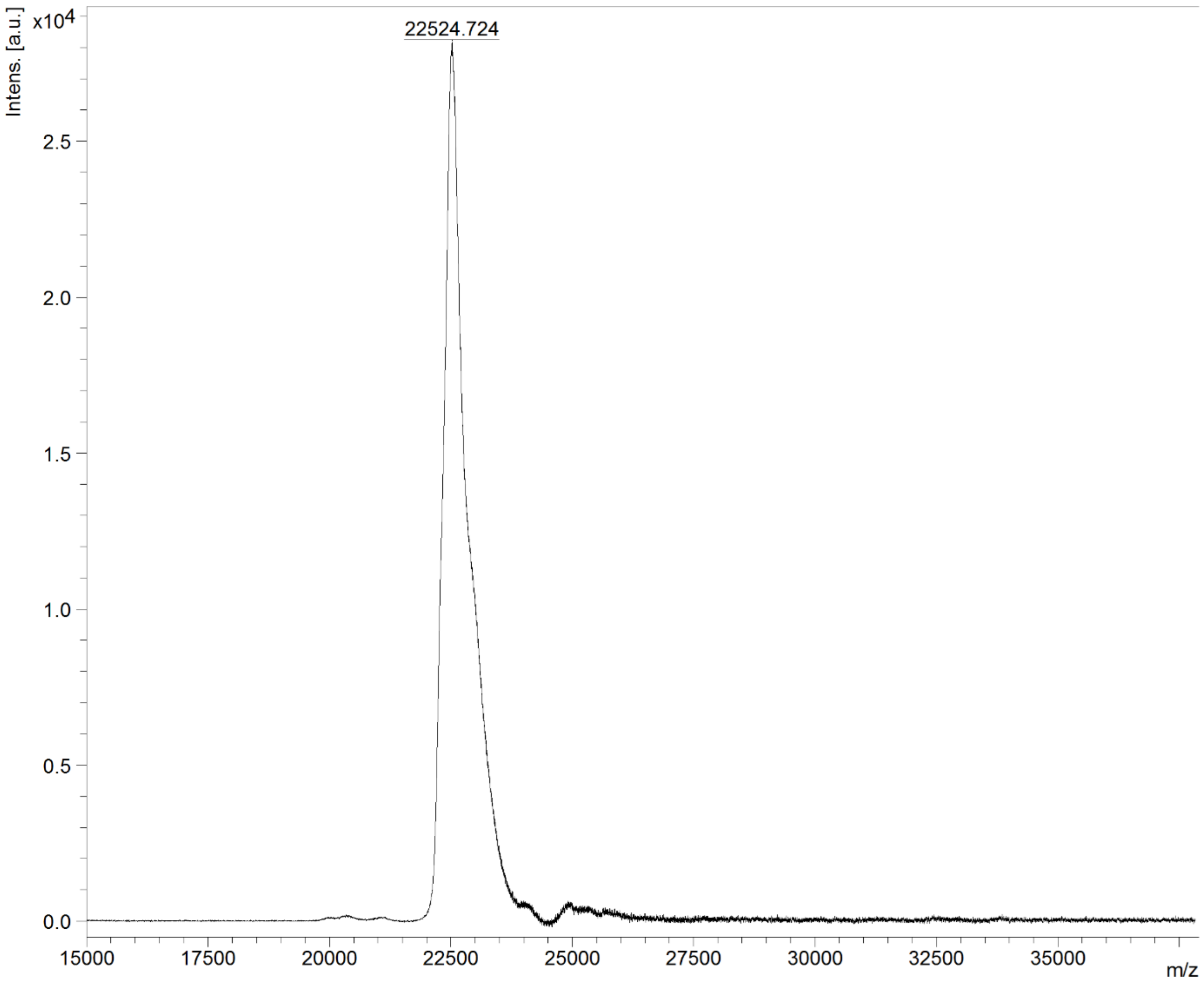


**Figure S19. MALDI-TOF Mass spectrum of purified CTPF-ELP in the range of m/z 15000 to 35000.** There is a prominent m/z peak at 22524.724, which is close to the theoretically predicted Mw of CTPF-TEV 22607.638 kDa. Also, the absence of the PhoCl1 empty barrel peak determines its purity.

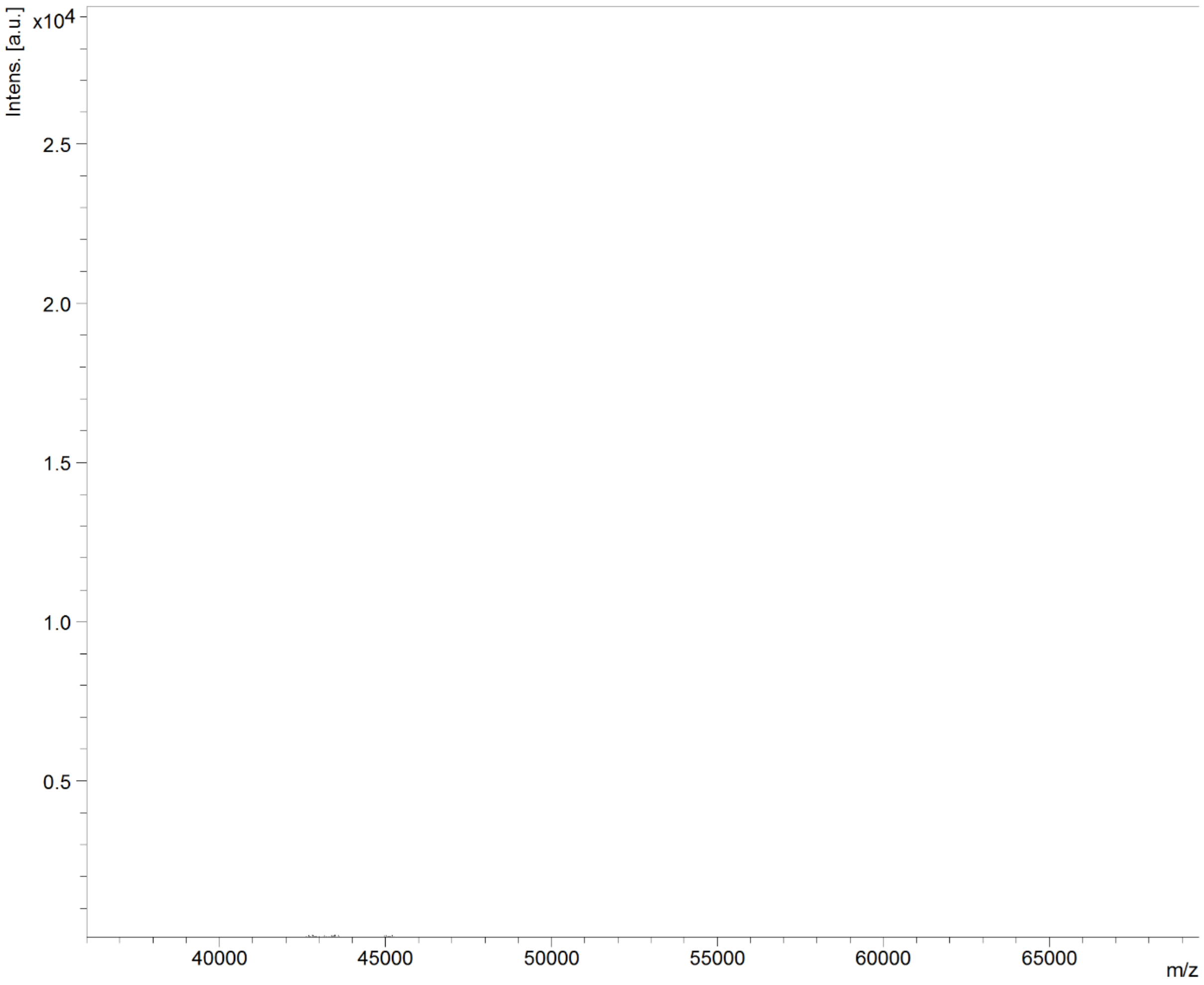


**Figure S20. MALDI-TOF Mass spectrum of purified CTPF-ELP in the range of m/z 40000-65000.** Absence of any significant peak in this region indicates the purity of CTPF-ELP and discards the possibility of fusion protein contamination.

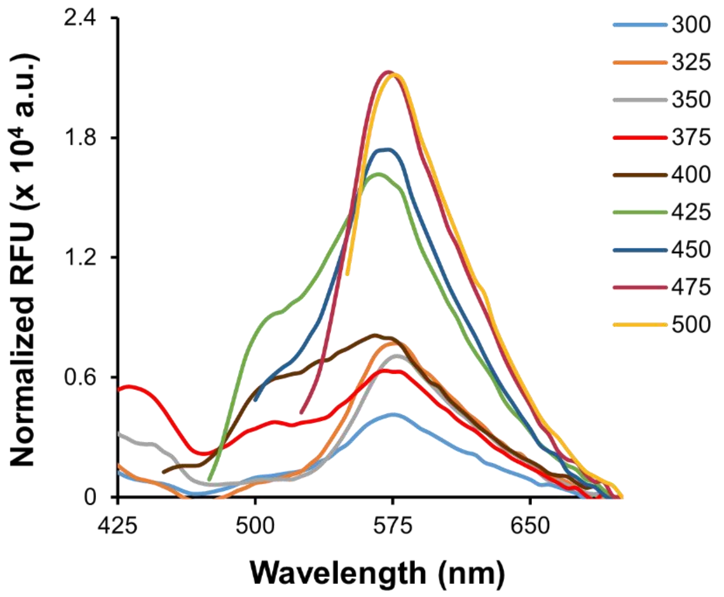


**Figure S21.** Fluorescence spectrum of CTPF at different excitation wavelengths.

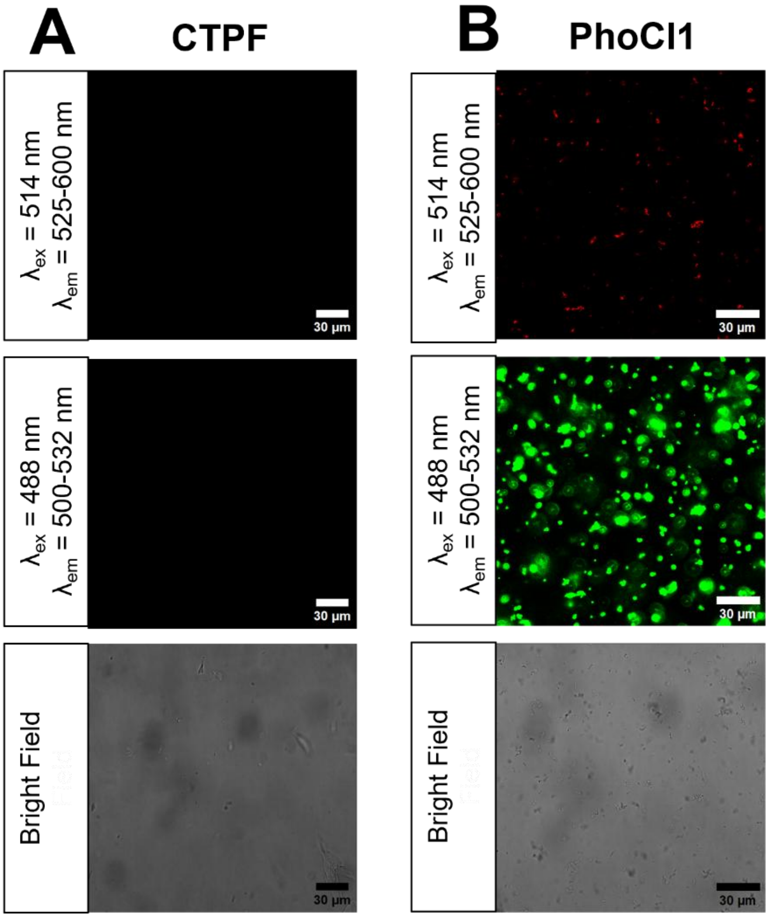


**Figure S22. Fluorescence microscopy of CTPF and PhoCl1 in the presence of PEG. (A)** CTPF shows no phase separation upon the addition of PEG and **(B)** The PhoCl1 shows the presence of fluorescence in the green channel and red channel, but no overlaps, differentiating it from CTPF-ELP coacervates.

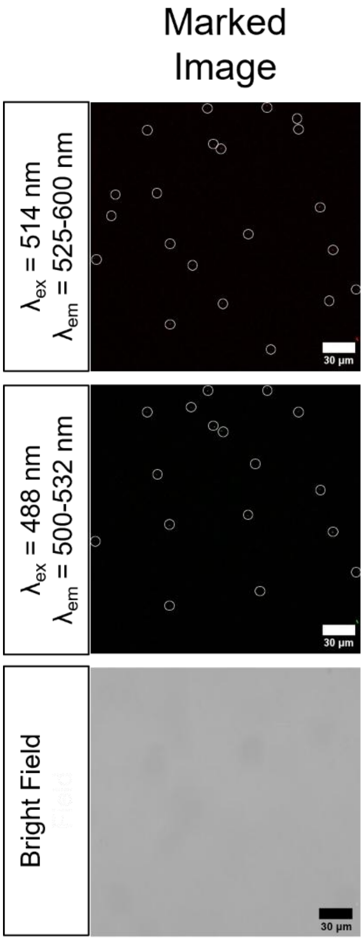


**Figure S23. Tagging of R5 with CTPF.** Confocal images of CTPF-R5 observed under the red and green channels along with bright field micrographs. Circular markings around the coacervates have been provided to guide the reader to the coacervate.

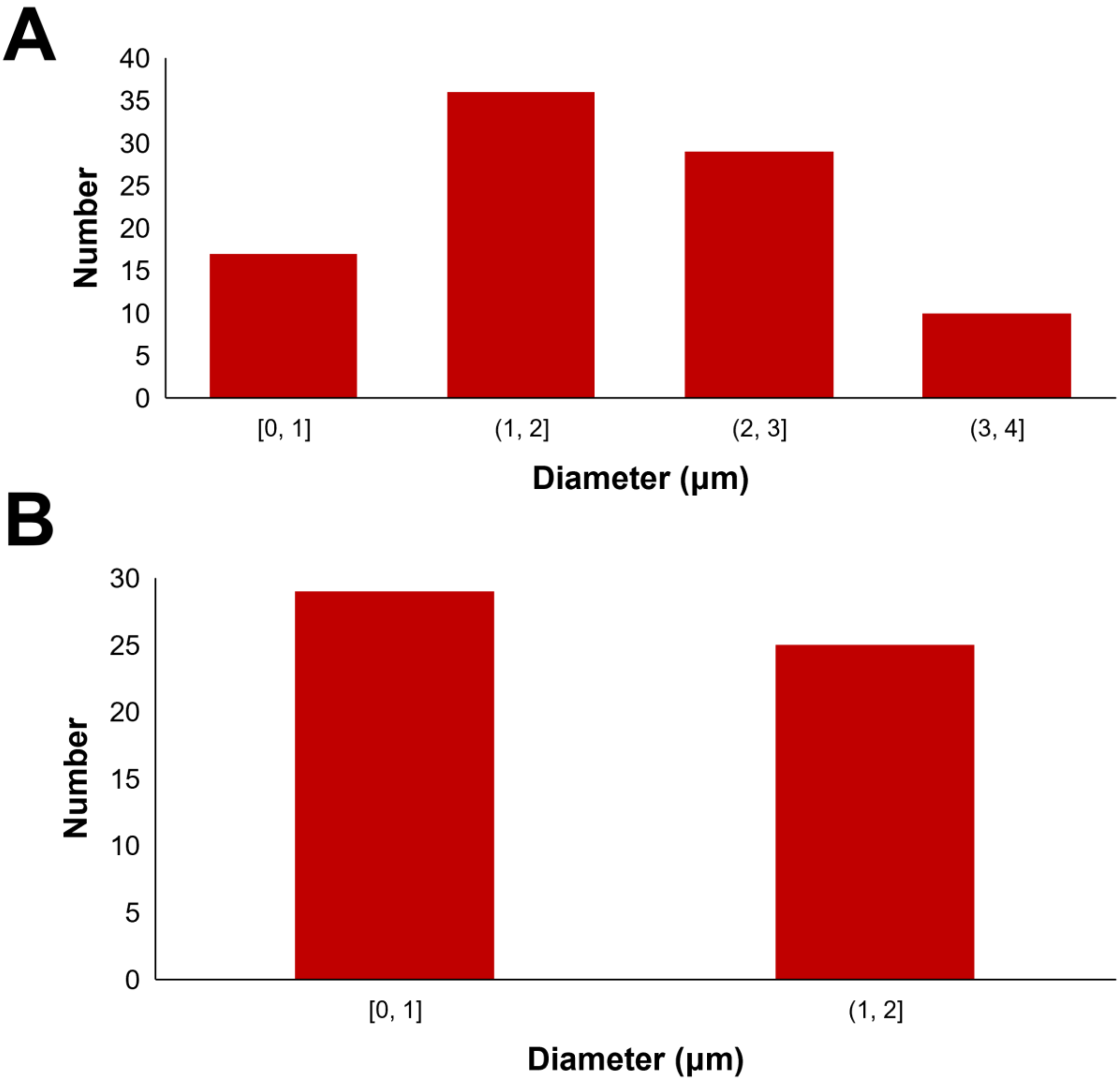


**Figure S24. Size distribution of the coacervates. (A)** Average size of CTPF-ELP coacervates is calculated to be 2.24 ± 0.9 μm (Coacervate Number: 92) and **(B)** Average size of CTPF-R5 coacervates is calculated to be 0.93 ± 0.3 μm (Coacervate Number: 54).
